# Paired vagus nerve stimulation drives precise remyelination and motor recovery after myelin loss

**DOI:** 10.1101/2024.05.10.593609

**Authors:** Rongchen Huang, Elise R. Carter, Ethan G. Hughes, Cristin G. Welle

## Abstract

**Summary:** Myelin loss in the central nervous system can cause permanent motor or cognitive deficits in patients with multiple sclerosis (MS). While current immunotherapy treatments decrease the frequency of demyelinating episodes, they do not promote myelin repair or functional recovery. Vagus nerve stimulation (VNS) is a neuromodulation therapy which enhances neuroplasticity and the recovery of motor function after stroke, but its effects on myelin repair are not known. To determine if VNS influences myelin repair, we applied VNS following a demyelinating injury and measured longitudinal myelin dynamics and functional recovery. We found that VNS promotes remyelination by increasing the generation of myelinating oligodendrocytes. Pairing VNS with a skilled reach task leads to the regeneration of myelin sheaths on previously myelinated axon segments, enhancing the restoration of the original pattern of myelination. Moreover, the magnitude of sheath pattern restoration correlates with long-term motor functional improvement. Together, these results suggest that recovery of the myelin sheath pattern is a key factor in the restoration of motor function following myelin loss and identify paired VNS as a potential remyelination therapy to treat demyelinating diseases.

---

Oligodendrocytes are the myelinating cells in the central nervous system and regulate properties of neuronal function, including conduction velocity, action potential timing and population synchronization^1^. Loss of oligodendrocytes and their myelin sheaths in demyelinating disorders, such as multiple sclerosis (MS), leads to axonal dysfunction and subsequent cognitive and motor deficits^2,3^. Current immunotherapies to treat MS reduce the frequency of relapse, but do not promote the restoration of lost myelin or prevent the accumulation of disability^4^. Remyelination restores conduction velocity and promotes neurophysiological recovery^5,6^, underscoring the therapeutic potential of myelin repair. Remyelination therapies indicate promising early results^7,8^, highlighting a clear need for new treatments that can enhance remyelination and restore lost function.

Vagus nerve stimulation (VNS), a well-established neuromodulation therapy for the treatment of epilepsy and depression^9,10^, was recently approved by the FDA to treat motor impairment following stroke^11^. For motor recovery following neurologic injury, stimulation is paired with relevant motor tasks, leading to circuit-specific neural plasticity^11,12^. However, it is currently unknown if paired VNS can promote myelin repair or accelerate functional recovery following demyelinating injury.

The pattern of myelination is highly variable across different cell types and brain regions^13,14^. Such diversity in myelin patterns is crucial for fine-tuning axonal conduction velocity within neuronal networks, which is vital for precise behaviors like sound localization^15^. However, demyelinating injuries disrupt the precise pattern and the remyelination process often introduces myelin sheaths to previously unmyelinated locations^16–18^. Factors that influence the restoration of myelin pattern and whether myelin pattern is important to mediate functional recovery following demyelination remain unknown.

To examine whether VNS can promote myelin repair and drive functional improvement following a demyelinating injury, we tracked the dynamics of oligodendrocytes and individual myelin sheaths before, during, and after demyelinating injury using longitudinal *in vivo* two-photon imaging. We found that VNS promotes remyelination via enhancing the generation of new oligodendrocytes. Pairing VNS with a skilled reach task led to the regeneration of myelin sheaths on previously myelinated axon segments, enhancing the restoration of the original pattern of myelination. Paired VNS improved motor recovery through increased reach consistency producing a rapid consolidation of an expert reach trajectory. Moreover, the magnitude of restoration of the myelin pattern correlated with improved long-term motor performance. These results indicate a new role for paired VNS to enhance oligodendrogenesis, guide myelin sheath restoration, and enhance long-term functional improvement, suggesting that VNS has the potential to become an important remyelination therapy.

## Results

### VNS promotes remyelination via enhancing oligodendrogenesis after myelin injury

To visualize the dynamics of myelin loss and repair, we performed longitudinal two-photon *in vivo* imaging in the forelimb region of primary motor cortex in mice expressing enhanced green fluorescent protein (EGFP) in mature oligodendrocytes and myelin (*MOBP-EGFP* mice^19^). To induce demyelination, ten-week-old mice were fed 0.2% cuprizone diet for three weeks^16^. Two weeks after cuprizone treatment onset, mice were implanted with a stimulation cuff on the left cervical vagus nerve^20^ (Micro-Leads Inc) and head-mounted connector pins (**Fig. 1a**). Three days after removal of the cuprizone diet, mice received a daily VNS session for seven days (**Fig. 1b**, ‘VNS’). Stimulation pulse number and parameters were predetermined and based on our prior work (22.94±1.28 pulses)^12^; **Extended Data Fig. 1a**). The efficacy of VNS was confirmed during the surgical implantation by change in heart rate and respiration rate^20^ (**Extended Data Fig. 1d,e**; **Methods**).

To determine whether VNS can modulate recovery from demyelinating injury, we first confirmed that cuprizone treatment induced a significant loss of mature oligodendrocytes and their myelin sheaths following three weeks of treatment (**Fig. 1c,d**) consistent with our previous work^16^. Importantly, we found that the extent of cuprizone-mediated demyelination did not differ between the VNS and the unstimulated surgical sham control groups (**Fig. 1e**).Removal of cuprizone treatment resulted in robust oligodendrogenesis that correlated to the extent of cuprizone-mediated oligodendrocyte loss^16^ (also see **Extend Data Fig. 1b**). Therefore, we normalized the number of regenerated oligodendrocytes as a proportion of total oligodendrocytes loss (‘OL replacement’; see **Methods**). We found that oligodendrocyte replacement was ∼15% higher in VNS group following four weeks of recovery (Unstimulated: 55.36±4.32% versus VNS: 71.11±2.79%, **Fig. 1f**). To further characterize this regenerative oligodendrogenesis, we quantified oligodendrocyte replacement using a three-parameter (3P) Gompertz equation. We examined the inflection point (when the rate switches from accelerating to decelerating) and asymptote (the plateau) of regenerative oligodendrogenesis^16^. We found that the inflection point was delayed 5 days by VNS, indicating that VNS increases the rate of oligodendrogenesis for a longer duration compared to Unstimulated mice (Unstimulated: 6.14±0.68 days post cuprizone versus VNS: 11.46±0.64 days post-cuprizone; **Fig. 1g**). Furthermore, VNS increased the asymptote value of oligodendrocyte replacement to 92.27±3.30% compared to 58.64±2.38% in the Unstimulated group (**Fig. 1g**). We did not find differences in the rate of oligodendrocyte replacement at baseline, during stimulation, and post-stimulation phases (**Extend Data Fig. 1c**), however, we found the replacement rate was increased in the VNS compared to Unstimulated group at the first week post-stimulation (**Fig. 1h**). Yet, across the entire post-stimulation period, the maximum rate of oligodendrocyte replacement did not differ between groups (**Fig. 1i**). Together, these data indicate that VNS enhances remyelination via increasing oligodendrogenesis during the first week after stimulation.

**Figure 1.**
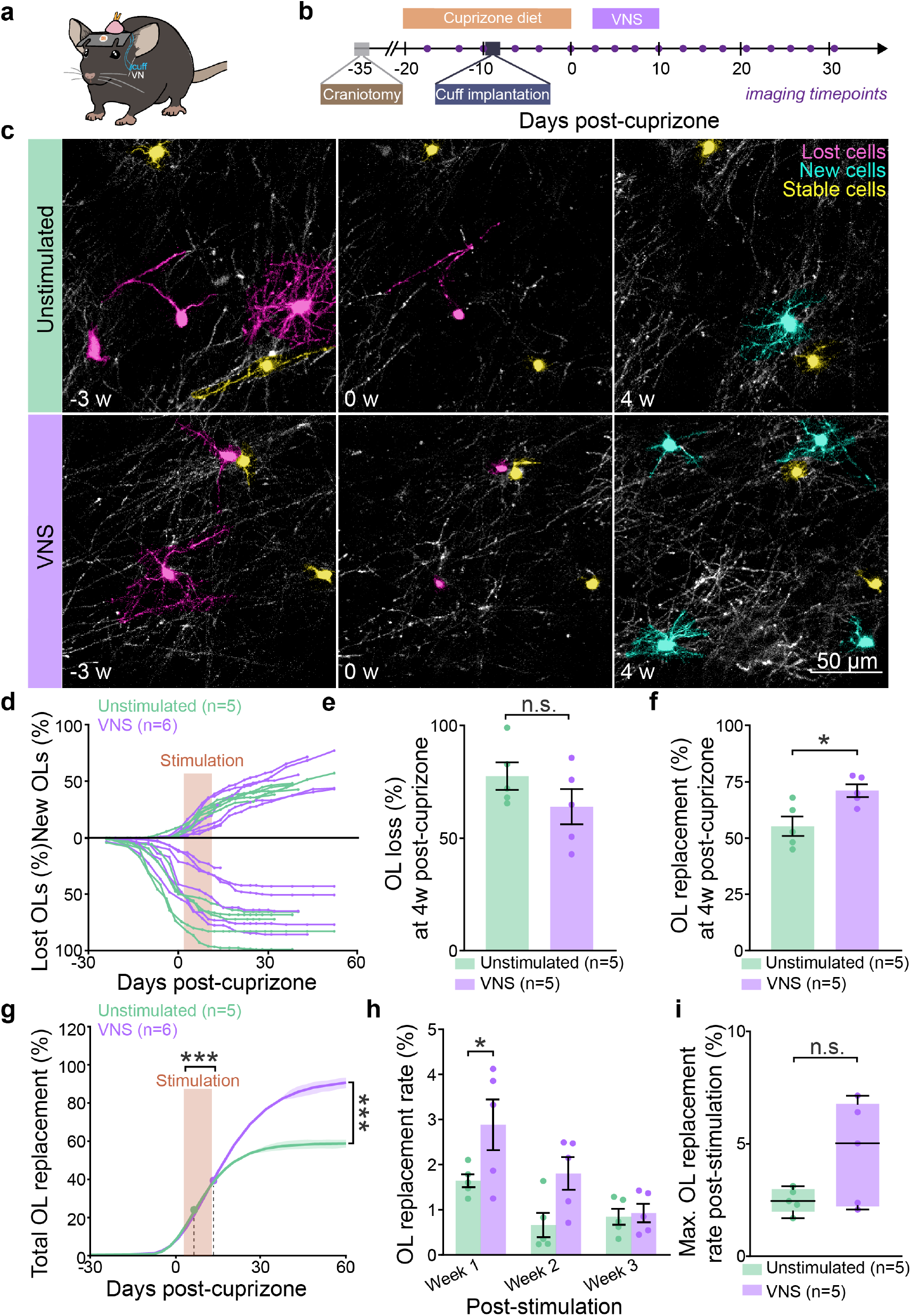
VNS enhances oligodendrogenesis after cuprizone-mediated demyelination. **a**, Mice are implanted with cranial imaging window and headbar and cuff around vagus nerve (VN). **b**, Experimental timeline. **c**, Oligodendrocytes are lost (pink), stable (yellow) or newly generated (blue) over a 7 weeks of imaging. 0 wk indicates the removal of the cuprizone diet. **d**, Cumulative OL gain and loss relative to baseline (percentage); Each trace represents individual mouse, rectangle indicates days with VNS. **e**, The percentage of OL loss at 4 weeks post-cuprizone is not different between groups (Student’s t-test; p=0.21). **f**, VNS increases the percentage of OL replacement at 4 weeks post-cuprizone (Student’s t-test; p=0.015). **g**, Gompertz 3P modeling for cumulative percentage of OL replacement shows that VNS increases asymptote value and inflection point, indicated by dot (Asymptote value: Student’s t-test; p<0.0001; Inflection point: Student’s t-test; p=0.0002). **h**, VNS increases OL replacement rate during the first week of post-stimulation (Two-way ANOVA with the Bonferroni correction t-test: week 1: p=0.034; week 2: p=0.056; week 3: p=0.93). **i**, The maximum OL replacement rate during post-stimulation phase is not different between two groups (Welch’s test; p=0.11). Lines and shaded areas in **g** represent the mean ± s.e.m. Boxplots in **i** represent minimum, median and maximum. *p<0.05, ***p<0.001, n.s., not significant; bars and error bars represent the mean ± s.e.m.

### VNS paired with a skilled reaching task enhances oligodendrocyte replacement

Previously, we showed that behavioral interventions promote remyelination following demyelinating injury^16^. VNS paired with skilled motor tasks enhances motor recovery following stroke and we recently showed that this intervention accelerates skilled motor learning in healthy animals^11,12,21^. We hypothesized that pairing VNS with skilled motor learning following demyelination would further accelerate oligodendrocyte restoration. To test this, we trained mice to perform a skilled, single-pellet forelimb reach task for seven days, with VNS delivered immediately following a reach success (‘Paired VNS’ versus ‘Motor Learning’ respectively) (**Fig. 2a,b**).

**Figure 2.**
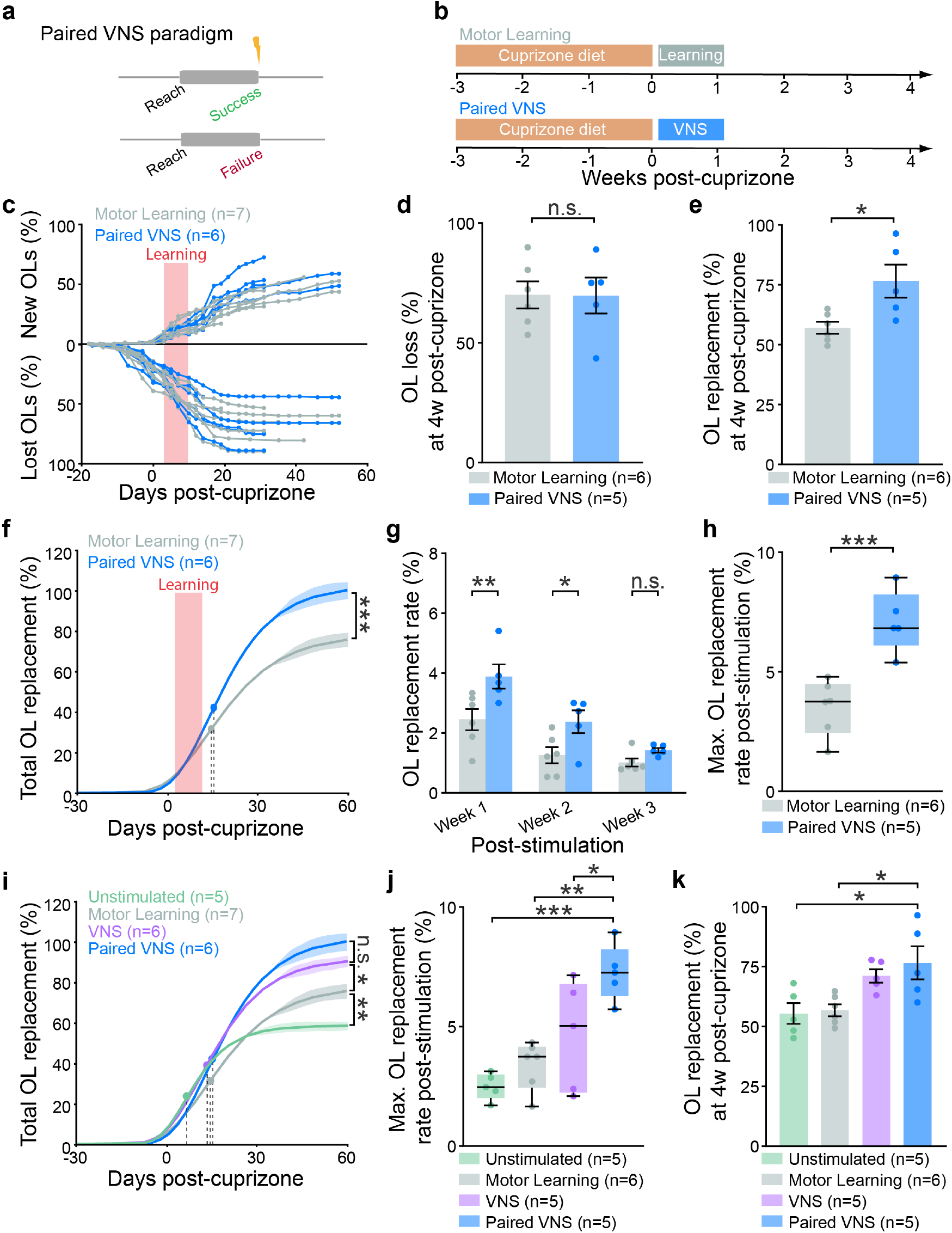
Pairing VNS with successful skilled reaching enhances oligodendrocyte replacement. **a**, Paired VNS paradigm. Stimulation is delivered on success outcomes but not failure outcomes. **b**, Experimental timeline with cuprizone and motor learning intervention. Motor Learning mice receive training task only, while Paired VNS receive motor learning with VNS being delivered on successful reaches. **c**, Cumulative OL gain and loss relative to baseline (percentage); Each trace represents an individual mouse, rectangle indicates days with motor learning. **d**, The percentage of OL loss at 4 weeks post-cuprizone is not different between groups (Student’s t-test; p=0.98). **e**, Paired VNS increases the percentage of OL replacement at 4 weeks post-cuprizone (Student’s t-test; p=0.019). **f**, Gompertz 3P modeling for cumulative percentage of OL replacement shows that Paired VNS increases asymptote value (Asymptote value: Student’s t-test; p=0.0008). **g**, Paired VNS increases OL replacement rate in the first two weeks of post-stimulation (Two-way ANOVA with Bonferroni correction t-test: week 1: p=0.0057; week 2: p=0.037; week 3: p>0.99). **h**, Paired VNS increases the maximum OL replacement rate during post-stimulation phase (Student’s t-test; p=0.0002). **i**, Gompertz 3P modeling for cumulative percentage of OL replacement shows that Motor Learning increases asymptote value compared to Unstimulated group, and both VNS and Paired VNS achieve an higher asymptote value compared to Motor Learning (One-way ANOVA with Tukey’s HSD: Unstimulated vs. Motor Learning, p=0.0066; Unstimulated vs. VNS, p=0.0002; Unstimulated vs. Paired VNS, p<0.0001; Motor Learning vs. VNS, p=0.024; Motor Learning vs. Paired VNS, p<0.0001; VNS vs. Paired VNS, p=0.22). **j**, Paired VNS increases the maximum OL replacement rate during post-stimulation phase compared to other three groups (One-way ANOVA with Tukey’s HSD, Unstimulated vs. Paired VNS, p=0.0003; Motor Learning vs. Paired VNS, p=0.0013; VNS vs. Paired VNS, p=0.036). **k**, Paired VNS causes the largest effects on OL replacement by 4 weeks post-cuprizone among four groups (One-way ANOVA with Tukey’s HSD: Unstimulated vs. Paired VNS, p=0.017; Motor Learning vs. Paired VNS, p=0.022; VNS vs. Paired VNS, p=0.82). Lines and shaded areas in **f&i** represent the mean ± s.e.m. Boxplots in **h&j** represent minimum, median and maximum *p<0.05, **p<0.01, ***p<0.001, n.s., not significant; bars and error bars represent the mean ± s.e.m.

First, we investigated the effects of Paired VNS on oligodendrogenesis in healthy mice. We found that Paired VNS altered the dynamics of oligodendrogenesis during learning (**Extend Data Fig. 2**). Previously, we found that motor learning transiently reduced the oligodendrogenesis rate during training^16^. Paired VNS eliminated the training-induced suppression compared to Motor Learning (Motor Learning: 2.20 ± 1.37% versus Paired VNS: 14.90±2.98% new oligodendrocytes, **Extend Data Fig. 2d,e**), although this did not change in the total number of new oligodendrocytes (Motor Learning: 14.40±2.06 versus Paired VNS: 12.25±1.11). These data indicate that Paired VNS modulates the dynamics of oligodendrogenesis in the healthy brain.

Next, we explored the influence of Paired VNS on oligodendrogenesis following demyelination injury (**Fig. 2b**). Following three weeks of cuprizone treatment, Paired VNS did not alter oligodendrocyte loss, consistent with results from the VNS condition (**Fig. 2c,d**). By four weeks following removal of cuprizone, Paired VNS increased the oligodendrocyte replacement by ∼20% (Motor Learning: 57.06±2.49% versus Paired VNS: 76.56±6.91%, **Fig. 2e**). While the inflection point did not differ between groups (Motor Learning: 12.88±0.91 days versus Paired VNS: 14.16±0.62 days), the asymptote value of oligodendrocyte replacement for Paired VNS mice increased compared to Motor Learning animals (Motor Learning: 77.45±3.75% versus Paired VNS: 102.17±3.58%; **Fig. 2f**). The rate of oligodendrocyte replacement was increased by Paired VNS in the first two weeks of post-stimulation phase compared to Motor Learning animals (**Fig. 2g**). Furthermore, Paired VNS increased the maximum rate of oligodendrocyte replacement compared to Motor Learning group (Motor Learning: 3.38±0.42% versus Paired VNS: 7.26±0.53%, **Fig. 2h**).

To examine if Paired VNS provides additional benefits on remyelination, we compared the oligodendrocyte replacement between Unstimulated, Motor Learning, VNS, and Paired VNS groups. Following cuprizone removal, Paired VNS increased the asymptote value of oligodendrocyte replacement compared to Unstimulated and Motor Learning groups but not to the VNS group (Unstimulated: 58.64 ± 2.38%; Motor Learning: 77.30 ± 3.75%; VNS: 92.27 ± 3.30%; Paired VNS: 102.17 ± 3.58%, **Fig. 2i**). Over the four weeks following demyelination, we found that Paired VNS increased the maximum oligodendrocyte replacement after stimulation to a greater extent than all other groups (Unstimulated: 2.49±0.25%; Motor Learning: 3.38±0.42%; VNS: 4.61±1.03%; Paired VNS: 7.26±0.53%, **Fig. 2j**). Also, Paired VNS increased oligodendrocyte replacement compared to Unstimulated and Motor Learning groups but did not differ compared to the VNS group (Unstimulated: 55.36±4.32%; Motor Learning: 57.06±2.49%; VNS: 71.11±2.79%; Paired VNS: 76.56±6.91%, **Fig. 2k**). Together, these data show that Paired VNS and VNS both increase regenerative oligodendrogenesis following demyelinating injury.

### Paired VNS promotes individual sheath replacement to previously myelinated locations

Remyelination after injury often alters the placement of myelin sheaths, and the corresponding myelin pattern^16–18^. To determine the effect of VNS on myelin sheath placement, we examined the number, the length, and the location of sheaths generated by new oligodendrocytes during remyelination (**Fig. 3a**). We found no changes in the number (**Fig. 3b**) nor the length (**Fig. 3c**) of individual sheaths generated by new oligodendrocytes across all groups. Next, we modeled the restoration of the sheath population using the mean number of sheaths per new oligodendrocyte per mouse (**Fig. 3b**). Four weeks following cuprizone removal, we found that the restoration of myelin sheaths in the VNS group did not differ from Unstimulated, Motor Learning, or Paired VNS groups. In contrast, Paired VNS increased the restoration of sheaths in compared to Unstimulated and Motor Learning groups, and was able to fully restore sheath number to original levels (Unstimulated: −30.36±6.47%; Motor Learning: - 26.72±3.82%; VNS: −10.08±3.44%; Paired VNS: −3.16±7.29%, **Fig. 3d**).

**Figure 3.**
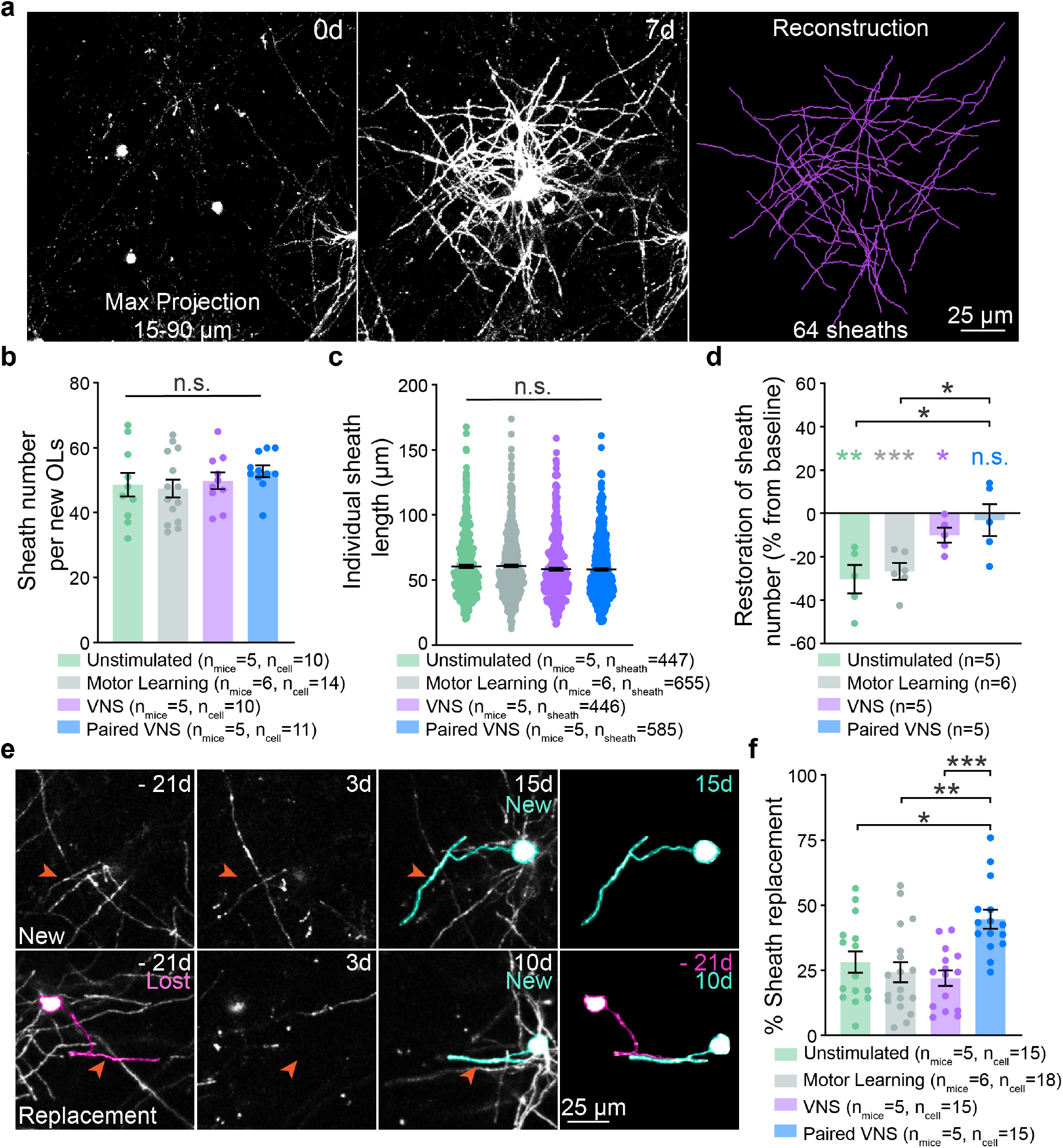
Paired VNS drives remyelination by regenerated myelin sheaths. **a**, Representative image of individual new oligodendrocyte and associated myelin sheaths. 0d indicates the day of cuprizone removal. **b**, The number of sheaths generated from individual new oligodendrocytes is similar among four groups (One-way ANOVA; p=0.55). **c**, Individual sheath length is similar among four groups (Kruskal-Wallis test; p=0.077). **d**, Paired VNS increases the percentage of restored sheaths compared to Unstimulated and Motor learning (One-way ANOVA with Tukey’s HSD; Unstimulated vs. Paired VNS, p=0.013; Motor Learning vs. Paired VNS, p=0.027; VNS vs. Paired VNS, p=0.81). Only Paired VNS fully restores lost myelin sheaths by 4 weeks post-cuprizone compared to baseline (One sample t test with control set as 0; p=0.0094, p=0.0009, p=0.043 and p=0.69, respectively). **e**, Representative images of a new sheath located at previously unmyelinated areas (‘New’, top) and a new sheath located at previously myelinated areas (‘Replacement’, bottom). **f**, Paired VNS increases the percentage of sheath replacement compared to other three groups (One-way ANOVA with Tukey’s HSD; Unstimulated vs. Paired VNS, p=0.012; Motor Learning vs. Paired VNS, p=0.0012; VNS vs. Paired VNS, p=0.0005). *p<0.05, **p<0.01, ***p<0.001, n.s., not significant; bars and error bars represent the mean ± s.e.m.

Following demyelination, myelin sheaths from regenerated oligodendrocytes are placed at previously unmyelinated locations (‘New’) or replaced at previously myelinated locations (‘Replacement’) (**Fig. 3e**). In line with previous work^16–18^, newly generated oligodendrocytes in the Unstimulated, Motor Learning, and VNS groups placed a majority of new sheaths in previously unmyelinated areas (**Extend Data Fig. 3a-c**). However, new oligodendrocytes in the Paired VNS condition regenerated new sheaths in new and replaced locations in equal proportions (‘New’; 54.43±3.63% of sheaths vs. ‘Replacement’; 44.58±3.63% of sheaths, **Extend Data Fig. 3d**). Across all groups, Paired VNS increased the replacement of sheaths at previously myelinated locations (Unstimulated: 28.16±4.16%; Motor Learning: 24.19±3.89%; VNS: 21.97±2.98%; Paired VNS: 44.58±3.63%, **Fig. 3f**). Taken together, these data suggest that Paired VNS restores myelin sheath number to pre-injury levels and enhances the replacement of sheaths to previously myelinated locations.

### Paired VNS restores myelin sheath pattern after demyelination

Paired VNS drives individual oligodendrocytes to place myelin sheaths onto previously myelinated axon locations, yet it remains unknown if these sheath-level differences accumulate into measurable effects on the pattern of myelination across a population of axons. To determine if Paired VNS influences the pattern of myelin regeneration, we examined restoration of the myelin sheath population by tracing all the sheaths within a 150 × 150 × 60 μm^3^ region at the onset of imaging (−21 days) and 31 days following removal of cuprizone (31 days). Consistent with the findings analyzing individual sheaths of new oligodendrocytes (**Fig. 3d**), Paired VNS restored the population of myelin sheaths to pre-injury conditions to a greater extent than Motor Learning (Motor Learning: −25.02±3.46% versus Paired VNS: - 1.79±6.09%; p=0.007, Student’s t-test).

To determine if regenerated sheaths were placed in previously myelinated locations, sheath locations were classified into one of three categories: non-restored (present before cuprizone-mediated demyelination but not afterwards), persistent (present before and following cuprizone-mediated demyelination), and de novo (not present before cuprizone-mediated demyelination) (**Fig. 4a-c**). To further characterize “persistent” sheaths, we reviewed the longitudinal image timeseries to determine if individual sheaths were either not lost during demyelination (‘Survived’) or were new sheaths restored onto previously myelinated locations (**Fig. 4d**). We found that Paired VNS increased the number of replaced sheaths, but did not alter the number of surviving sheaths compared to the Motor Learning group (**Fig. 4e**). Furthermore, while there was no difference in de novo sheaths between groups (Motor Learning: 20.20±1.96% versus Paired VNS: 24.65±2.36%; p = 0.18, Student’s t test), we found that Paired VNS increased the total persistent sheath number as compared with Motor Learning (**Fig. 4f,g**). Next, we quantified degree of similarity between the original and post-injury myelination pattern by examining the percentage of persistent sheaths of the total myelin sheath population before and after cuprizone-mediated demyelination. We found that Paired VNS increased myelin pattern similarity compared to the Motor Learning group (**Fig. 4h**). Together, these results indicate that Paired VNS promotes the placement of regenerated sheaths in previously myelinated locations, which drives the restoration of the original myelin sheath pattern.

**Figure 4.**
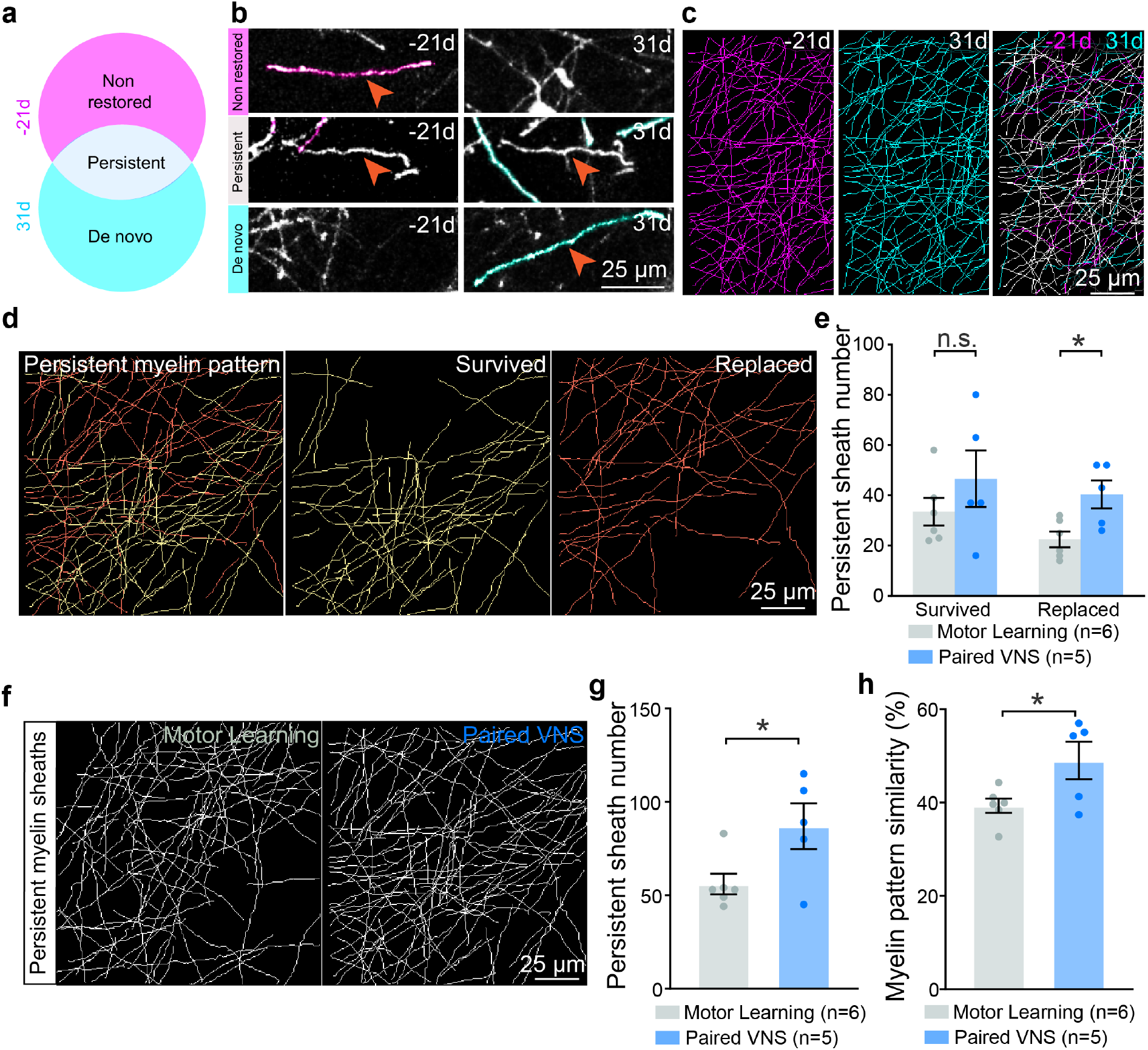
Paired VNS restores myelin pattern following demyelination. **a**, Venn diagram of three categories of sheaths based on their existence at 21 days before cuprizone only (‘Non restored’, magenta), 31 days post-cuprizone only (‘De novo’, cyan) or at both timepoints (‘Persistent’, grey). **b**, Representative images of a ‘Non restored’ (top), ‘Persistent’ (middle) and ‘De novo’ (bottom) sheath at 21 days before cuprizone (left column) and 31 days post-cuprizone (right column). **c**, Maximum projections of pseudo-colored tracings of all myelin sheaths within the same region at 21 days before cuprizone (left), 31 days post-cuprizone (middle) and an overlay of two timepoints (right). **d**, Example of a persistent myelin pattern (left) that contains sheaths that ‘survived’ across all imaging timepoints (middle, yellow) and regenerated sheaths that ‘replaced’ at previously myelinated locations (right, orange). **e**, Paired VNS increases the number of replaced sheaths, but does not change the number of survived sheaths (Multiple unpaired t-test; Survived: adjusted p=0.16; Replaced: adjusted p=0.017). **f**, Representative sheath reconstructions of persistent myelin pattern from Motor Learning and Paired VNS groups. **g**, Paired VNS increases the number of persistent sheaths (Student’s t-test; p=0.036). **h**, Paired VNS increases the percentage of myelin pattern restoration after myelin injury (Student’s t-test; p=0.039). *p<0.05, n.s., not significant; bars and error bars represent the mean ± s.e.m.

### Paired VNS improves motor recovery following demyelinating injury

VNS paired with physical rehabilitation following a neurologic injury improves functional recovery in models of stroke, spinal cord injury and peripheral nerve injury^22,23^. To understand if paired VNS can similarly improve motor recovery after myelin loss, mice were first trained to proficiency on the forelimb reach task for 7 days (**Fig. 5a, Extended Data Fig. 4a,b**), and then exposed to three weeks of cuprizone treatment to induce demyelination. On the third day following the removal of cuprizone, the animals engaged in daily motor recovery sessions for 15 days (**Fig. 5a**). To determine the effects of demyelination on motor performance, we compared proficiency in the last two training sessions during motor learning (days −29 to −28 post-cuprizone) to early recovery (days 3 to 5 post-cuprizone) in individual mice. In control animals, we found that demyelination impairs motor performance (“Motor Recovery” Learning: 35.14±4.48% vs. “Motor Recovery” Recovery 24.56±4.03%, **Fig. 5b**). Yet, paired VNS rescued motor performance to pre-injury levels in the early recovery period (“Motor Recovery” Recovery: 24.56±4.03% vs. “Paired VNS Recovery” Recovery: 44.41±4.99%, **Fig. 5b**) and led to lasting motor functional improvement across the 15 days of recovery (Motor Recovery: 29.17±4.17% vs. Paired VNS Recovery: 42.99±3.19%, **Fig. 5c**). In addition to mean performance, Paired VNS improved performance on the worst-performing day for each animal compared to Motor Recovery, indicating that Paired VNS increases motor performance stability across sessions (Motor Recovery: 16.87±4.22% vs. Paired VNS Recovery: 27.80±2.48%, **Fig. 5d**).

**Figure 5.**
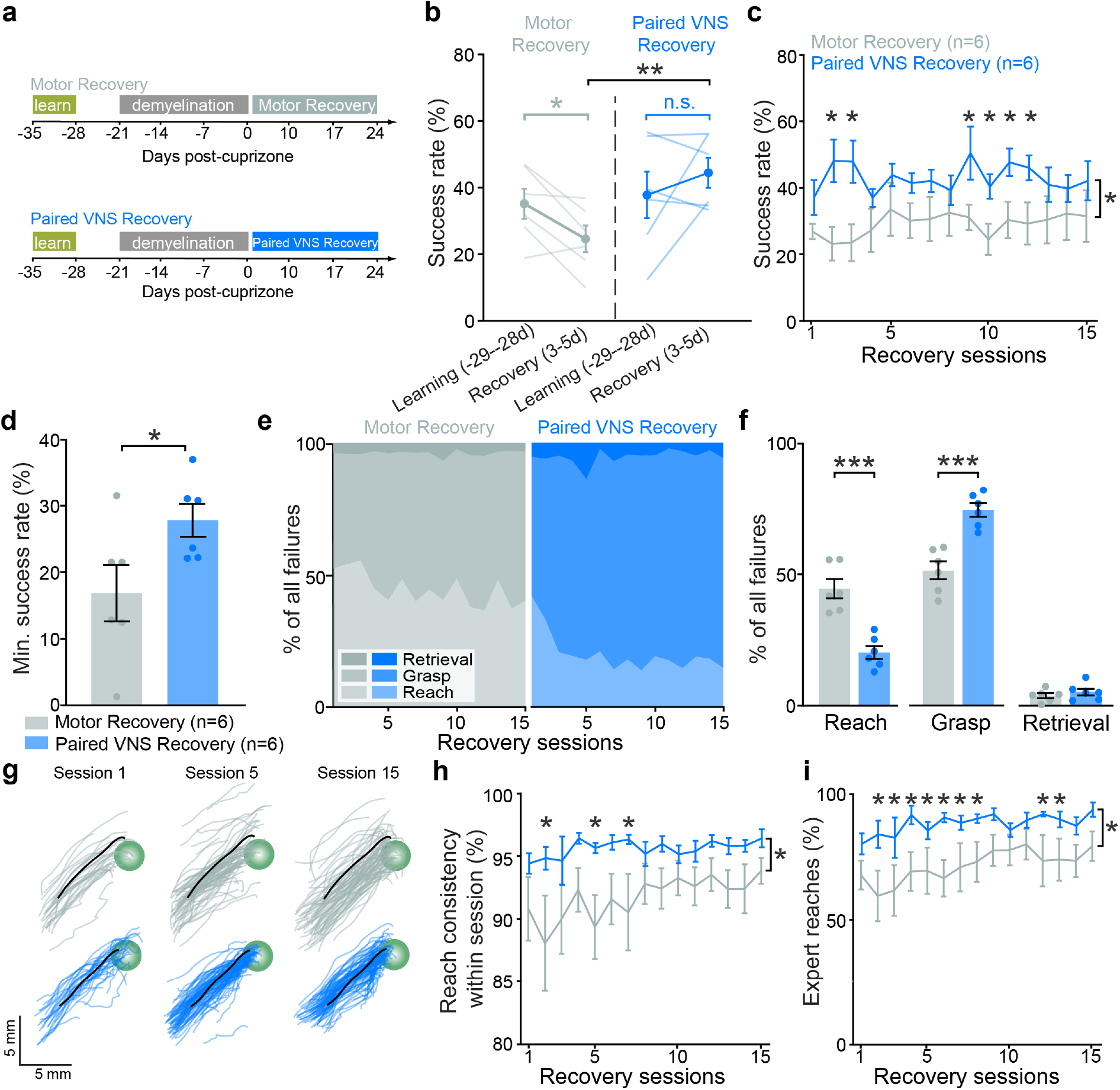
Paired VNS improves functional recovery after demyelination. **a**, Timeline to test the effects of paired VNS on functional recovery after demyelination. **b**, The mean success rate of the first three days post-cuprizone is significantly lower compared to their pre cuprizone level in Motor Recovery group (Paired t-test; t(5)=2.73, p=0.041), while the mean success rate of the first three days post-cuprizone is similar compared to their pre cuprizone level in Paired VNS Recovery group (Paired t-test; t(5)=1.02, p=0.36). The mean success rate of the first three days post-cuprizone is significantly higher in Paired VNS Recovery group compared to Motor Recovery mice (Student’s t-test; t(10)=3.29, p=0.0081). **c**, During recovery phase, Paired VNS Recovery mice performed significantly better over Motor Recovery mice (REML with Benjamini and Hochberg post hoc (false discovery rate(FDR)= 10%); F(1)= 5.89, p=0.036). **d**, Paired VNS increases the minimum success rate during recovery phase compared to Motor Recovery mice (Student’s t-test; t(10)=2.33, p= 0.049). **e**, Breakdown of failure reaches for each group over 15 recovery sessions. Light colors: reach failure; intermediate: grasp failure; dark: retrieval failure. **f**, A comparison of different types of failure outcomes between two groups. Paired VNS decreases reach failure (Student’s t-test; t(10)=5.49, p= 0.0003), and increases grasp failure compared to Motor Recovery group (Student’s t-test; t(10)=5.35, p= 0.0003). **g**, Examples outward trajectories from a Motor Recovery mouse (top) and a Paired VNS Recovery mouse (bottom) at different sessions. Black lines represent each mouse’s “expert reach.” Green circle represents the location of food pellet. **h**, Paired VNS increases the reach consistency within each training session (REML with Benjamini and Hochberg post hoc (false discovery rate(FDR)= 10%); F(1)= 5.58, p=0.040). **i**, Paired VNS increases the percentage of expert reaches over training sessions (REML with Benjamini and Hochberg post hoc (false discovery rate(FDR)= 10%); F(1)= 5.47, p=0.041). *p<0.05, **p<0.01, ***p<0.001, n.s., not significant; bars and error bars represent the mean ± s.e.m.

To understand how Paired VNS improves motor performance, we explored the features of individual reaches across the recovery sessions. First, we classified failed reaches into failure of hand targeting to the pellet (“Reach”), failure of pellet grasp (“Grasp”) and failure of pellet retrieval (“Retrieval”) (**Fig. 5e**). Paired VNS reduced the targeting failures, while increasing grasp failures as compared to Motor Recovery (‘Reach’: Motor Recovery: 44.62±3.68% vs. Paired VNS Recovery: 20.31±2.46%; ‘Grasp’: Motor Recovery: 51.52±3.40% vs. Paired VNS Recovery: 74.45±2.62%, **Fig. 5f**), indicating that Paired VNS improves reach accuracy. To explore how Paired VNS may influence reach accuracy, we tracked the three-dimensional trajectory of the reach using deep learning methods^12,24,25^. Paired VNS did not alter reach velocity or path length compared to the Motor Recovery group (**Extended Data Fig. 4c,d**). Instead, Paired VNS increased the similarity of individual reach trajectories within a training session (**Fig. 5g**), resulting in increased pair-wise correlations within session (“Reach consistency”), starting in early recovery (Motor Recovery: 91.74±1.58% vs. Paired VNS Recovery: 95.64±0.46%, **Fig. 5h**). We next examined if the increased reach consistency was influenced by the animal’s expert reach trajectory, defined as the average successful reach trajectory from the last two days of recovery (sessions 14-15). Paired VNS increased the percentage of expert reaches (reaches within 95% similarity of the expert reach trajectory) across all training sessions, with effects beginning in early recovery (Motor Recovery: 71.59±6.85% vs. Paired VNS Recovery: 88.34±2.10%, **Fig. 5i**) before paired VNS-induced alterations to oligodendrocyte replacement (3d post cuprizone, Motor Learning: 9.50± 1.45% versus Paired VNS: 10.41±3.47%; **Fig. 2f**). We found that this effect was driven by increased similarity of failure reaches to the expert reach trajectory (Motor Recovery: 93.86±1.22% vs. Paired VNS Recovery: 96.84±0.39%, **Extended Data Fig. 4e,f**), again indicating improved motor consistency in Paired VNS Recovery mice. Together, these data demonstrate that paired VNS improves motor recovery through improved reach consistency with the expert trajectory very early in the recovery process, preceding myelin repair processes.

### Paired VNS drives long-term motor learning

While Paired VNS enhances motor learning in healthy mice^12^, myelin loss impairs motor learning^16^. To test if Paired VNS enhances motor learning after demyelinating injury, we examined motor learning immediately following cuprizone-mediated demyelination. We found that Paired VNS did not affect training proficiency compared to the Motor Learning group (Motor Learning: 32.79. ± 2.58 %; Paired VNS: 34.96 ± 2.72 %, p=0.57; Student’s t test), indicating that Paired VNS does not rescue motor learning immediately following demyelination.

In healthy mice, experience-dependent adult oligodendrogenesis is required for memory consolidation and retrieval^26,27^. To determine whether enhanced remyelination by Paired VNS results in long-term functional improvements, we retrained mice on the forelimb reach task 11 weeks after cuprizone (10 weeks after motor learning) (**Fig. 6a**). Importantly, during the seven-day retrieval protocol, neither Paired VNS nor Motor Learning groups received VNS. While mice in the Motor Learning group did not improve over retrieval (**Fig. 6b,c**), we found that Paired VNS improved their performance across the retrieval sessions (**Fig. 6d,e**). Together these data show that while Paired VNS does not alter learning following demyelination, this intervention leads to long-term functional improvements likely by influencing reach-related motor circuits.

**Figure 6.**
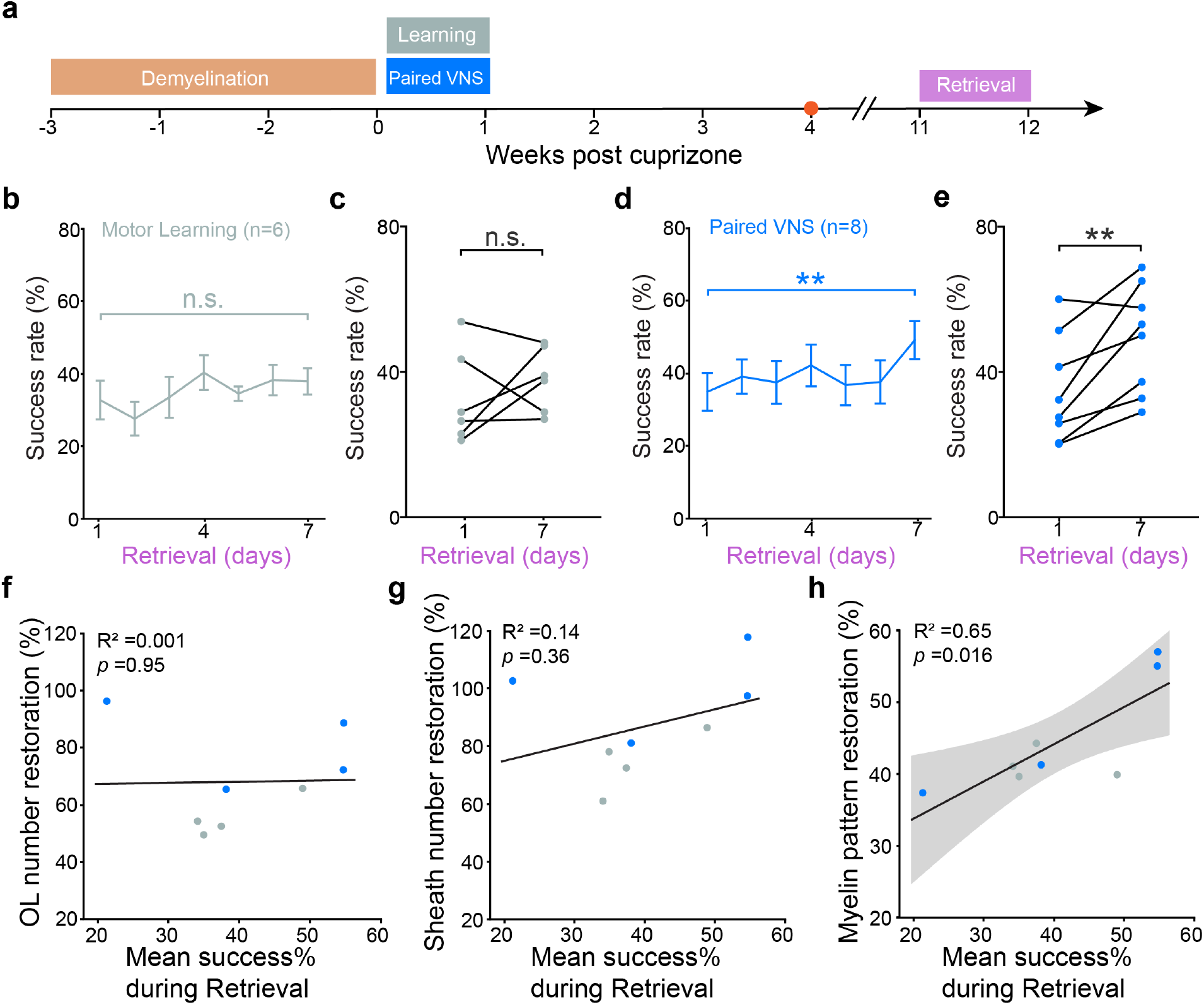
Paired VNS drives long-term functional improvement of motor performance. **a**, Experimental timeline with retrieval intervention. Imaging obtained at 4 weeks post-cuprizone (red dot) was used to measure different remyelination features: OL number restoration (**f**), sheath number restoration (**g**), and myelin pattern restoration (**h**). **b**, Motor Learning mice do not show an improvement in motor performance during seven-day retrieval phase (REML; p=0.10). **c**, Success rate is similar between day 1 and day 7 in Motor Learning group (Paired t-test; p=0.42). **d**, Paired VNS mice show an improvement in motor performance during seven-day retrieval phase (REML; p=0.0087). **e**, Success rate is significantly higher at day 7 compared to day 1 in Paired VNS group (Paired t-test; p=0.0086). **f-h**, Mean success% during retrieval phase is not correlated with the magnitude of OL number restoration (**f**) non sheath number restoration (**g**) but is highly correlated with the magnitude of myelin pattern restoration (**h**) (standard least squares regression, shading represents 95% CI). **p<0.01, n.s., not significant; bars and error bars represent the mean ± s.e.m.

In the healthy brain, learning drives changes to the pattern of myelination of behaviorally-relevant axons which are correlated with enhanced behavioral performance^28^. Distinct myelin patterns along individual axons are critical for precise spike timing within neuronal circuits, which is essential for temporal processing in sound localization^15^. Whether restoration of myelin pattern following demyelinating injury promotes long-term functional recovery remains unknown. We explored whether the long-term functional improvements were linked to the extent of remyelination for individual mice. We found that the number of restored oligodendrocytes at four weeks following the removal of cuprizone was not correlated to motor performance during retrieval (**Fig. 6f**). Similarly, we found no correlation between the number of restored sheaths and functional improvement on the reach task (**Fig. 6g**). However, we found that restoration of the myelin pattern was strongly correlated with behavioral proficiency during retrieval (**Fig. 6h**). Together, these data indicate that Paired VNS drives long-term functional improvement in behavioral performance on a fine-skilled motor task through enhanced restoration of the original myelin pattern in forelimb motor cortex.

## Discussion

New therapeutic approaches are needed to regenerate oligodendrocytes, replace myelin sheaths, and restore normal function following myelin loss. In this study, we establish that VNS promotes remyelination following a demyelinating episode via enhancing oligodendrogenesis. When paired with skilled reach learning, we find that VNS restores the original myelin pattern by enhancing the number and placement of new sheaths on axon locations that were myelinated prior to injury. Furthermore, Paired VNS improves motor recovery and promotes long-term retention of motor learning. Importantly, the degree of myelin pattern restoration correlates to long-term motor performance, suggesting that the preserved myelin pattern promotes restoration of motor behavior. These results propose a novel role for VNS therapy to drive myelin repair following a demyelinating episode, restoring myelin pattern and leading to improved functional outcomes.

### VNS enhances oligodendrogenesis following myelin loss

Myelin plasticity, or the dynamic process of oligodendrocyte addition and myelin sheath remodeling, plays an important role in cognition and motor learning. Generation of new oligodendrocytes in adulthood is required for motor learning and consolidation of new memories^26,27,29,30^. Following myelin injury, pharmacological enhancement of oligodendrogenesis alleviates motor deficits and reverses cognitive dysfunction^31,32^. Similarly, precisely-timed motor learning significantly enhances oligodendrogenesis in motor cortex after demyelination^16^. Here, we find that VNS doubles the rate of oligodendrocyte regeneration over motor learning alone, effectively restoring myelin content to pre-injury levels (**Fig. 3d**). Interestingly, VNS-enhanced oligodendrogenesis is similar for VNS with motor learning and VNS applied alone without concomitant motor activity (**Fig. 2i,k**). Previously, VNS has been shown to potently drive widespread cortical activity^12,33^ and neuronal activity modulates oligodendrocyte generation^34^, suggesting that neural activity could be a key factor leading to VNS-enhanced oligodendrogenesis.

### Paired VNS leads to specific sheath replacement and myelin pattern restoration

The anatomical variation in the pattern of myelination tunes conduction velocity and the precise timing of action potentials to optimize temporal processing across brain regions^35–37^. Global demyelination leads to an incomplete myelin repair and a dramatic reorganization of the pattern of myelination^16,17^. Along individual axons, endogenous remyelination restored the overall myelin content but not the precise pattern of myelination, leading to the hypothesis that opportunistic behavior of oligodendrocytes defines the remyelination probability of individual axons^13^. However in contrast to newly generated oligodendrocytes, surviving oligodendrocytes preferentially place sheaths at previously myelinated locations^16,38^, suggesting that the precision of remyelination may amenable to modulation. Our data demonstrate that paired VNS increases myelin sheath placement in previously myelinated locations and restores the original pattern of myelination (**Fig. 3e and 4h**). These findings raise the possibility that beyond increasing the total myelin content of the remyelination response via pharmacological enhancement^7,8^, additional cues to guide the specificity of remyelination may further improve functional recovery following demyelinating injury.

The mechanisms governing sheath placement following demyelinating injury are not well understood. While previous studies found that components of the node of Ranvier are rapidly dissembled following demyelination^39^, recent work shows that key components remain in place potentially providing a spatial template for remyelination^17^. Recent studies using single-oligodendrocyte demyelination find evidence both in support and against the reestablishment of node of Ranvier locations^18,40^, pointing to the need for investigation with large-scale demyelination events. Alternatively, neuronal activity may modulate the regeneration of myelin patterning. In healthy animals, chemogenetic activation drives preferential myelination of activated axons^41^ and motor learning can alter pattern of myelination on behaviorally-relevant axons^28^. Following a demyelinating episode, optogenetic activation drives oligodendrogenesis, but does not produce preferential myelination on activated axons in the white matter^42^. Our data suggest that Paired VNS leads to short- and long-term changes in motor performance, potentially first optimizing motor circuit activity during early recovery leading to activity-dependent myelin patterning which results in long-term functional improvements.

### Long-term functional recovery with paired VNS

Electrical stimulation of the vagus nerve paired with sensorimotor input confers potent and specific neural plasticity. In healthy animals, when paired with a sensory stimuli or movement, VNS enhances map plasticity of the corresponding primary cortical area^43,44^ and improves motor learning^12^. VNS paired with physical rehabilitation following a neurologic injury improves functional recovery in models of stroke, spinal cord injury and peripheral nerve injury^22,23^. Similarly, following a demyelinating injury, Paired VNS improved functional recovery through increased reach consolidation towards the expert reach (**Fig. 5i**). By refining motor behavior, Paired VNS may modulate task-relevant subsets of neurons, as seen during motor learning in healthy animals^12^, driving activity-dependent myelination^28^. In contrast, paired VNS did not influence the rate of motor learning following myelin loss, likely due to the difficulty in learning the skilled reach behavior in injured animals and the high degree of variability across individuals. However, the animals that received paired VNS during learning demonstrated improved the long-term motor performance during the retrieval session at 11 weeks post-injury (**Fig. 6d,e**). Furthermore, the amount of long-term functional improvement correlated to the magnitude of myelin pattern restoration. This implies that circuit-specific neural activation could be a key factor leading to VNS-enhanced myelin pattern restoration. We propose that following demyelinating injury, Paired VNS leads to restoration of motor circuits needed for motor performance through circuit-specific myelination. This is consistent with the known role for myelination in the formation of remote, but not early, memories in fear conditioning and spatial navigation tasks^26,27^, and points towards a role for myelination pattern in long-term memory formation.

### Neuromodulatory systems in VNS-driven myelin repair

The therapeutic effects of VNS on the treatment of epilepsy and depression depend, at least in part, on activation of the noradrenergic locus coeruleus^45,46^, and Paired VNS enhances motor learning through phasic activity in the cholinergic basal forebrain^12^. Patients with MS have reduced norepinephrine (NE) in cortex^47^, and the use of a NE reuptake inhibitor alleviates symptoms^48^. In healthy animals, arousal-induced NE release enhances calcium dynamics in oligodendrocyte precursor cells (OPCs)^49^ and chemogenetic activation of locus coeruleus neurons promotes differentiation of oligodendrocyte precursor cells (OPCs) into myelinating oligodendrocytes^50^. Following cuprizone-induced demyelination, the administration of an acetylcholinesterase (AchE) inhibitor accelerates remyelination by enhancing differentiation potentially through the activation of nicotinic acetylcholine (Ach) receptors^51^, but in contrast, Ach activation of muscarine 1 receptors inhibits OPC differentiation^31,52^. Together, this presents VNS activation of cholinergic or noradrenergic neuromodulatory systems as a possible mechanism for enhanced oligodendrogenesis following myelin loss.

### The anti-inflammatory effects of VNS

MS, an autoimmune disease, is associated with activation of immune cells. Current immunotherapies targeting effector T cells and B cells reduce relapse rates and slow disease progression by mitigating the inflammatory response^53^. The vagus nerve is a key component of the autonomic nervous system, and VNS has shown promise in reducing inflammation in diseases like rheumatoid arthritis (RA) and inflammatory bowel disease (IBD)^54,55^. VNS modulates the vagus nerve-mediated inflammation reflex through cholinergic signaling pathway via α7 nicotinic acetylcholine receptors on splenic macrophages to decrease the release of circulating pro-inflammatory cytokines^56–58^. In this study, we utilized the cuprizone-mediated demyelination model to determine the effects of VNS on regenerative capacity without the confound of autoimmunity. The cuprizone model has limited blood-brain barrier damage and minimal infiltration of peripheral immune cells into the central nervous system^59^. Future work is needed to determine if the potential anti-inflammatory effects of VNS could work in synergy with the remyelination effects to provide a potent therapeutic option for patients with multiple sclerosis.

## Materials and Methods

### Animal care

All animal experiments were conducted in accordance with protocols approved by the Animal Care and Use Committee at the University of Colorado Anschutz Medical Campus. Male and female mice used in these experiments were kept on a 14-h light/ 10-h dark schedule with ad libitum access to food and water, aside from training-related food restriction (see the section “**Forelimb reach training with CLARA system**”). All mice were randomly assigned to conditions and were age-matched (±7 days) across experimental groups. C57BL/6 MOBP– EGFP lines, which have been previously described (*17, 20*), were used for two-photon imaging and behavioral training. C57BL/6 wildtype mice were used for behavioral training. Mice were group housed before cuff implantation surgery and single-housed following surgery and throughout behavior training.

### Surgeries

#### Cranial window surgery

Six- to eight-week-old mice were anesthetized with isoflurane (induction, 5%; maintenance, 1.0–2.0%, mixed with 0.6 liter per minute of O_2_) and kept at 36.5 °C body temperature with a thermostat-controlled heating plate. Local application of 1% injectable lidocaine was performed at incision sites, and eye ointment was applied to prevent corneal drying. A 2 × 2 mm^2^ cranial window (0–2 mm anterior to bregma and 0.5–2.5 mm lateral to midline) was created above the primary motor cortex on the left cerebral hemisphere using a high-speed dental drill. A piece of cover glass (VWR, No. 1) was then placed in the craniotomy and sealed with GLUture (WPI). A custom metal plate with a central hole was then attached to the skull with dental cement (C&B Metabond), allowing for future *in vivo* imaging. After the headplate was secured, mice received intraperitoneal meloxicam (5 mg/kg) following surgery to manage pain and were then returned to home cage with heating pad beneath it until they woke up.

#### Vagus nerve stimulating cuff implantation

Cuff implantation surgeries were performed as previously described^12,20^. Commercial cuffs from Micro-Leads (150 μm cuffs) were used for all the surgeries. These cuffs were soldered to gold pins and soaked in ethanol for 24 hrs before use. Three months old mice with cranial windows were anesthetized with isoflurane and maintained similarly as described above. The vagus nerve was accessed by making an incision around the ventral cervical region, and the nerve was dissected from the surrounding carotid sheath. The cuff was tunneled subcutaneously to the ventral cervical incision from an incision made at the connection of the dental cement and the skull. The vagus nerve was placed inside the cuff. The ventral cervical incision was sutured using 6-0 absorbable sutures. Electrical connectors were fixed to the skull using dental cement. The efficacy of stimulation was measured using peripheral biomarkers including change in breathing rate and heart rate immediately after surgery^20^. Breathing rate changes were observed by the experimenter with naked eye, while heart rate changes were determined with a paw sensor (Mouse Stat Jr., Kent Scientific). Mice in both ‘VNS’ and ‘Paired VNS’ groups exhibited significant changes in both breathing rate and heart rate were included for data analysis. Mice that only exhibited changes in heart rate but not breathing rate were categorized as ‘VNS low ΔBR’, and they did not enhance remyelination with VNS (**Extended Data Fig. 1**). Following surgery, mice received subcutaneous lactated ringers (∼100 mL as necessary) and intraperitoneal meloxicam (5 mg/kg). Mice continued to receive intramuscular gentamicin (5 mg/kg) for 3 days to prevent infection. Stimulation was delivered at least 7 days after cuff implantation surgery. To minimize potential surgical effects on myelination, both the ‘Unstimulated’ and ‘Motor Learning’ groups underwent the same surgical procedure, with the exception that no cuff was implanted around the nerve.

### Vagus nerve stimulation experiment

Mice were assigned into experimental groups in cohorts of 6-11 mice at a time, interleaved to achieve a similar number of mice in each group. Experimenters were not blinded to mice’s designations. In the VNS group, mice received manual stimulation at pseudorandom intervals to achieve between 10-35 stimulations within a 20-mins session per day. The number of stimulation pulses VNS group received each day was determined based on our previous study^12^ (**Extended Data Fig. 1**), and also matched the stimulation number for Paired VNS group in this study. For the Paired VNS group, mice were stimulated following every successful trial. VNS was delivered through gold pins as a 500 ms train of 15 pulses, with a 100 μs phase duration at 30 Hz. Current amplitudes ranged from 0.4 to 0.6 mA, based on whether mice exhibited adverse effects (e.g. gasping, constantly scratching around neck) during stimulation. Stimulation parameters were controlled and delivered using PulsePal, which was then connected to a stimulation isolation unit (A-M Systems, Model 2200 Analog Stimulus Isolator) to control amperage.

### Cuprizone-mediated demyelination

Cortical demyelination was induced using 0.2% cuprizone (Sigma Chemical, C9012) in either C57BL/6 MOBP–EGFP mice or wildtype mice. The cuprizone was stored in a glass desiccator at 4 °C and mixed into powdered chow (Harlan). Mice were provided with cuprizone chow in custom feeders designed to minimize exposure to moisture, and they had ad libitum access to this diet for a duration of 3 weeks. Feeders were refilled three times a week, and fresh cuprizone chow was prepared weekly. Cages were changed weekly to avoid build-up of old cuprizone chow in bedding, and to minimize reuptake of cuprizone chow following cessation of diet via coprophagia. Because cuprizone intake was voluntary, there was variation in the maximum oligodendrocyte loss, ranging from 40% to 100%. Given the strong correlation between oligodendrocyte gain and oligodendrocyte loss (**Extended Data Fig. 1**), directly using oligodendrocytes gain value could skew our results. To control for variations in total oligodendrocyte loss and its subsequent effects on oligodendrocyte gain, we therefore used oligodendrocytes replacement to measure oligodendrocyte gain relative to the severity of loss with the following equation^16^:

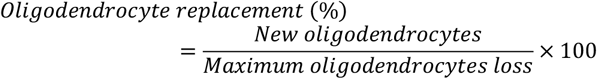

### Two-photon microscopy

*In vivo* imaging sessions began 2–3 weeks after the craniotomy and occurred 1–3 times per week for up to ten weeks, depending on the clarity of individual cranial windows. During imaging sessions, mice were anesthetized with isoflurane and immobilized by attaching the headplate to a custom stage. Images were taken using a Zeiss LSM 7MP microscope equipped with a BiG GaAsP detector and a mode-locked Ti:sapphire laser (Coherent Ultra) tuned to 920 nm. The average power at the sample during imaging ranged from 5 to 30 mW. Vascular and cellular landmarks were used to identify the same cortical area over longitudinal imaging sessions. Image stacks were acquired using a Zeiss W plan-apochromat ×20/1.0 NA water immersion objective giving a volume of 425 μm × 425 μm × 336 μm (1,024 × 1,024 pixels; corresponding to layers I–III, 0–336 μm from the meninges) from the cortical surface. On days involving motor tasks, images were collected after behavioral training to minimize potential effects of anesthesia on motor performance.

### Forelimb reach training with CLARA system

Motor training was conducted with CLARA system as previously described^12,24^. Prior to training, mice were weighed and habituated to CLARA training box for 20 mins one day before training initiation. During habituation, they were provided with three 20 mg food pellets (BIO-serv) near the slot of the training box where reaching behavior occurs. Following habituation, mice were food deprived and maintained at 85-90% of their baseline weight throughout training to sustain reach motivation. On day 1, mice were required to have three successful reaches before CLARA training session started. None of the groups received stimulation during these initial success trials. Both learning and retrieval sessions occurred for 7 consecutive days, with only one session per day, each lasting for 20 minutes. Recovery sessions lasted for three weeks, with five days a week. For all motor behavioral sessions, mice were required to reach for the pellet using their right forepaw.

High speed (150 Hz) video data was recorded from three FLIR Blackfly S cameras (model BFS-U3-16S2M-CS, Edmund Optics) placed at three angles (in front of the box, laterally to the box, and at a 45° angle above the box from the opposite side of the lateral camera). A neural network was trained for CLARA using manually annotated frames of the skilled reach behavior, labeling the hand center and the pellet. Video frames from all cameras were processed through this network in real-time to identify the location and state of the hand and pellet. This information was used to initiate trials through pellet placement and to categorize attempt outcomes as either ‘success’ or ‘failure’. Each trial began with the automated dispenser placing a food pellet on the post. The CLARA system determined the success or failure of the trial based on whether the mouse successfully retrieved the food pellet or knocked it off. If the outcome was a success, stimulation was delivered in a closed-loop manner. The timing of pellet placement, success or failure outcomes, and VNS delivery was recorded through CLARA.

### Behavioral and kinematic analysis

#### Behavioral curator analysis

Recorded videos from CLARA training sessions were processed overnight using custom Python scripts, with key reach timepoints being extracted: ReachInit, when the paw leaves the training box; ReachMax, the outward point of maximum distance from reach initiation; ReachEnd, when the hand returns to the cage. After processing, all reach trials were extracted with only those in which the food pellet was on the post being considered. The trial outcomes were post-hoc categorized into two main groups: ‘success’, where the mouse successfully took the pellet and transported it inside the box, and ‘failure’, which includes instances where the mouse failed to target the pellet (‘reach failure’), grasp the pellet (‘grasp failure’), or transport it inside the box (‘retrieval failure’). Mice that did not achieve success in at least 10% of their reach attempts throughout the entire training sessions were excluded from the analysis. To control for any batch effects on the forelimb reach training results, each batch included both control and experimental groups.

#### Kinematic analysis

The center of the paw and pellet were tracked in three-dimensional space during reach attempts. Tracking data was extracted using custom MATLAB scripts and documented as 3D arrays for kinematic analysis. Positional data points with <90% tracking confidence were replaced with interpolated values using a shape-preserving piecewise cubic Hermite interpolating polynomial (pchip). If interpolated data points comprised more than 50% of the data in a single reach, that reach was excluded from further kinematic analysis. Reach velocity was calculated as the 2-norm of Euclidean velocity and averaged over the period from ReachInit to ReachMax. To correct for tracking jumps that occur despite high confidence levels (i.e. to the left paw), positional data points with an associated absolute velocity exceeding 1000 mm/sec were also replaced with interpolated values. The path length was determined by the distance from ReachInit to ReachMax, calculated by performing numerical integration on the pchip data.

#### Reach consistency and expert reach

Positional data between ReachInit and ReachMax, referred to as trajectories, were normalized such that the pellet center was 0,0,0 and temporally warped as previously described^12,60^. The reach consistency within a session was calculated by averaging the pairwise correlation of all trajectories in a session. For each mouse, the expert reach trajectory was defined as the mean of all successful reach trajectories (ReachInit to ReachMax) collected on the final two days of training during the recovery phase. Individual trajectories were correlated with the expert reach trajectory, and reaches with >95% correlation are designated as an expert reach, and used to calculate the percent of expert reaches in a training session^12^.

### Imaging analysis

#### Imaging processing

Image stacks and time-series were analyzed using FIJI/ImageJ. All analyses were performed on unprocessed images except for sheath analysis, which were preprocessed with a Gaussian blur filter (radius = 0.8 pixel) to denoise and aid identification of individual myelin sheaths and nodes. When generating figures, maximum projections were used, and image brightness and contrast levels were adjusted for clarity. Longitudinal image stacks were registered using FIJI plugins ‘PoorMan3DReg’ and ‘Correct 3D drift’.

#### Cell tracking and cell number analysis

Custom Fiji scripts were used to track individual oligodendrocytes in four dimensions. This was achieved by identifying EGFP+ cell bodies at each time point, recording their x, y and z coordinates, and categorizing cellular behavior as either ‘new’, ‘lost’ or ‘stable’ cells. Oligodendrocyte gain and loss were cumulatively quantified relative to their own baseline cell number to account for variations in the baseline cell number. The rate of oligodendrocyte replacement was calculated as the percentage change in replacement over the amount of time elapsed. The maximum rate of oligodendrocyte replacement was determined as the highest value among all the calculated rates of oligodendrocyte replacement between two successive timepoints. For comparisons at 4 weeks post cuprizone mark, 18 mice used imaging timepoint at 31 days post-cuprizone, 2 mice (1 from Unstimulated group and 1 from VNS group) used imaging timepoint at 32 days post-cuprizone and 1 mouse (Motor Learning group) used imaging timepoint at 35 days post-cuprizone.

#### Myelin sheath analysis

*In vivo* z stacks were obtained from MOBP–EGFP mice using two-photon microscopy. These z stacks were processed with a 0.8-pixel Gaussian blur filter to aid in the identification of myelin internodes. Myelin paranodes and nodes of Ranvier were identified as previously described^16,19^, with paranodes exhibiting increased fluorescence intensity and nodes showing decreased to zero EGFP fluorescence intensity. To quantify sheath number and individual sheath length, myelin sheaths from newborn oligodendrocytes were traced using Simple Neurite Tracer at baseline (−21 days post-cuprizone), cuprizone withdrawal (0 day post-cuprizone), and the first timepoint of newborn oligodendrocyte being generated. Isolated GFP+ oligodendrocytes were randomly chosen for this analysis. Given that oligodendrocytes generate all sheaths in a limited period ^16^, only GFP+ myelin sheaths that was visible at the same time as oligodendrocyte cell body and in close proximity (less than ∼50 μm) to this cell body were considered connecting to that new oligodendrocyte.

To quantify sheath replacement of individual new oligodendrocytes, z stacks from baseline (−21 days post-cuprizone), cuprizone withdrawal (0 day post-cuprizone) and 4 weeks post-cuprizone (31-35 days post-cuprizone) were used. The same criterion as mentioned above was applied to determine whether the sheaths originated from the same newborn oligodendrocyte. To investigate the effects from VNS, only oligodendrocytes that were born after the application of VNS (that is from 3 to 31-35 days post-cuprizone) were chosen for this analysis. To quantify myelin pattern restoration, all myelin sheaths within a volume of 150×150×60 μm^3^ were traced using Simple Neurite Tracer at the baseline or 4 weeks post-cuprizone timepoint. Based on their presence at these two timepoints, they were categorized into three groups: ‘Non restored’ (present at baseline, but not at 4 weeks post-cuprizone); ‘Persistent’ (exhibiting more than 50% overlap between baseline and 4 weeks post-cuprizone); ‘De novo’ (present at 4 weeks post-cuprizone, but not baseline). To determine the subtype of a persistent sheath (‘survived’ or ‘replaced’), the sheath was examined across all imaging timepoints. A sheath was categorized as a ‘survived’ sheath if it existed at all imaging timepoints; otherwise, it was categorized as a ‘replaced’ sheath.

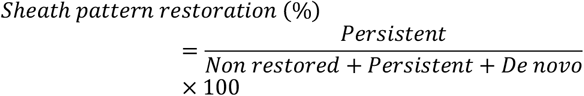

### Quantification and statistical analysis

A detailed and complete report of all statistics used, including definitions of individual measures, summary values, sample sizes and all test results, is provided in **Supplementary Table 1**. Statistical analyses were conducted using JMP (SAS) and GraphPad. Sample sizes were comparable to relevant publications^12,16,19^. All mice within a litter that underwent cranial window surgery, cuff implantation surgery, concurrent two-photon imaging and training timelines were considered as a batch. When possible, experimental groups were replicated in multiple batches with multiple experimental groups per batch. Normality test was performed for all datasets with the Shapiro-Wilk test. Nonparametric tests were used when normality was violated. Otherwise, parametric statistics, including paired and unpaired two-tailed Student’s t-tests (depending on the within- or between-subjects nature of the analysis), as well as analysis of variance (ANOVA) with appropriate post-hoc tests were performed. Two-tailed tests and α cutoff of < 0.05 were used unless otherwise specified. For statistical mixed modeling, we used a restricted maximum likelihood model (REML). All models were conducted with either one or two fixed effects, with ‘Mouse ID’ serving as a random variable, nested within batch if data came from different batches. Sigmoidal curves bound between 0 (baseline) and an estimated plateau were fitted to oligodendrocyte accumulation (either loss or replacement) across time using the 3P Gompertz equation modeling. R-squared and F-tests were computed for the regression models. In cases where significant effects were observed, the effect size was subsequently calculated using appropriate methods (**Supplementary Table 1**). For data visualization, error bars were used to represent the standard error of the mean (s.e.m), unless otherwise specified.

## Acknowledgments

We thank Clara Bacmeister for discussions on retrieval experiment; Spencer Bowles and Ryan Williamson for the troubleshooting on CLARA system; Kimberly Gagnon and Gustavo Della Flora Nunes for inputs on writing; all other members from Welle and Hughes for discussions.

## Author contributions

Conceptualization: EGH, CGW

Methodology: RH, EC, EGH, CGW

Investigation: RH

Visualization: RH, EGH, CGW

Funding acquisition: EGH, CGW

Project administration: EGH, CGW

Supervision: EGH, CGW

Writing – original draft: RH

Writing – review & editing: RH, EGH, CGW

## Data and materials availability

All data and materials that support the findings, tools, and reagents will be shared on an unrestricted basis; requests should be directed to the corresponding authors.

## EXTENDED DATA

**Extended Data Fig. 1.**
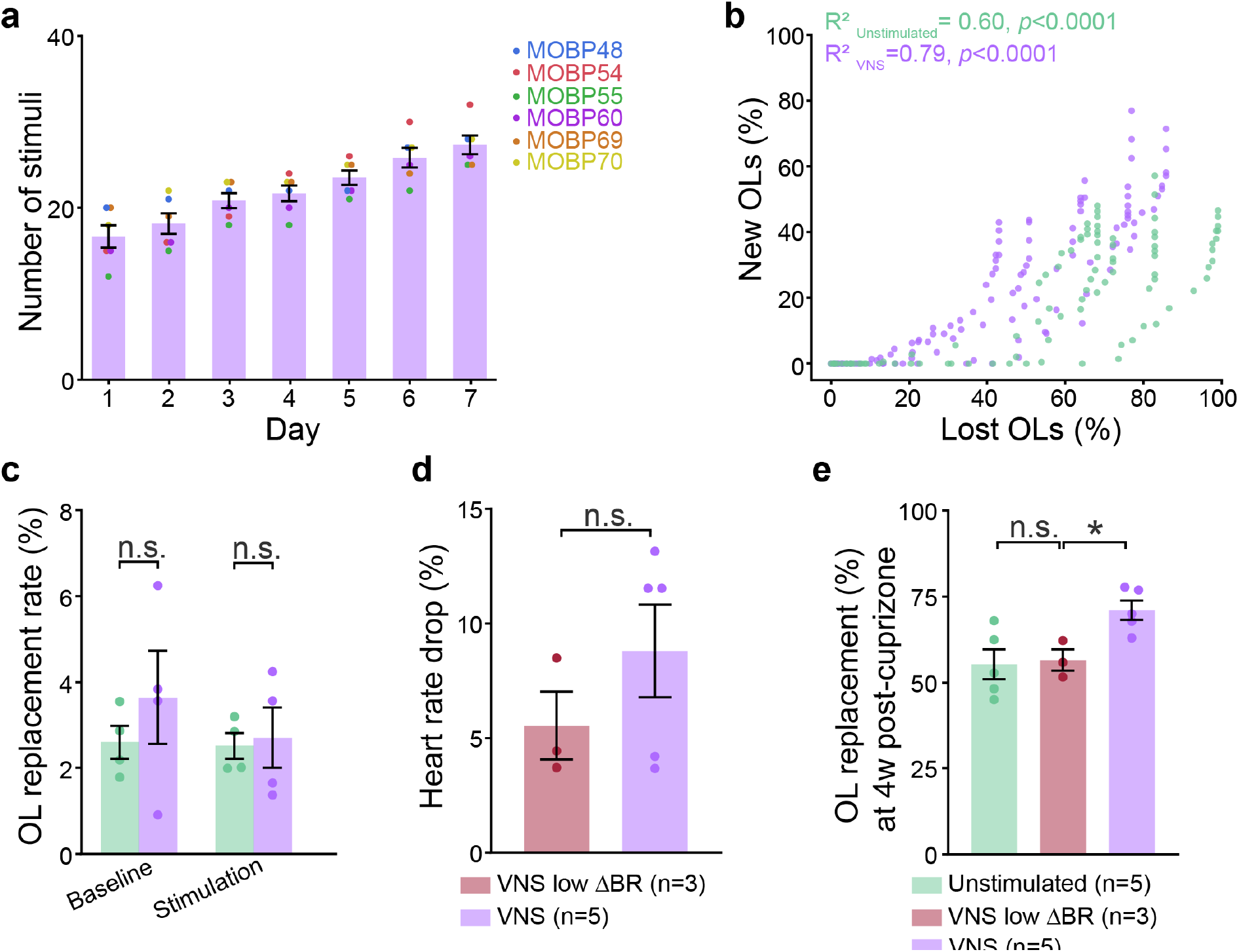
The efficacy of VNS is confirmed with respiration rate change and heart rate change. **a**, Number of stimuli each mouse in VNS group received across seven days. **b**, Cumulative OL gain is tightly correlated with OL loss for both Unstimulated and VNS groups (Spearman’s correlation). **c**, OL replacement rate during baseline and stimulation phases are not different between groups (Two-way ANOVA with Bonferroni post hoc: Baseline: p=0.62; Stimulation: p>0.99). **d**, The heart rate change tested during cuff implantation surgery is not different between VNS low ΔBR (breath rate change; details see Methods) and VNS groups (Student’s t-test; p=0.30). **e**, VNS low ΔBR group does not improve the percentage of OL replacement by 4 weeks post-cuprizone as compared to Unstimulated mice, and this value is significantly lower compared to VNS group (One-way ANOVA with Dunnett post-hoc test: Unstimulated vs. VNS low ΔBR: p=0.96; VNS low ΔBR vs. VNS: p=0.046). *p<0.05, n.s., not significant; bars and error bars represent the mean ± s.e.m.

**Extended Data Fig. 2.**
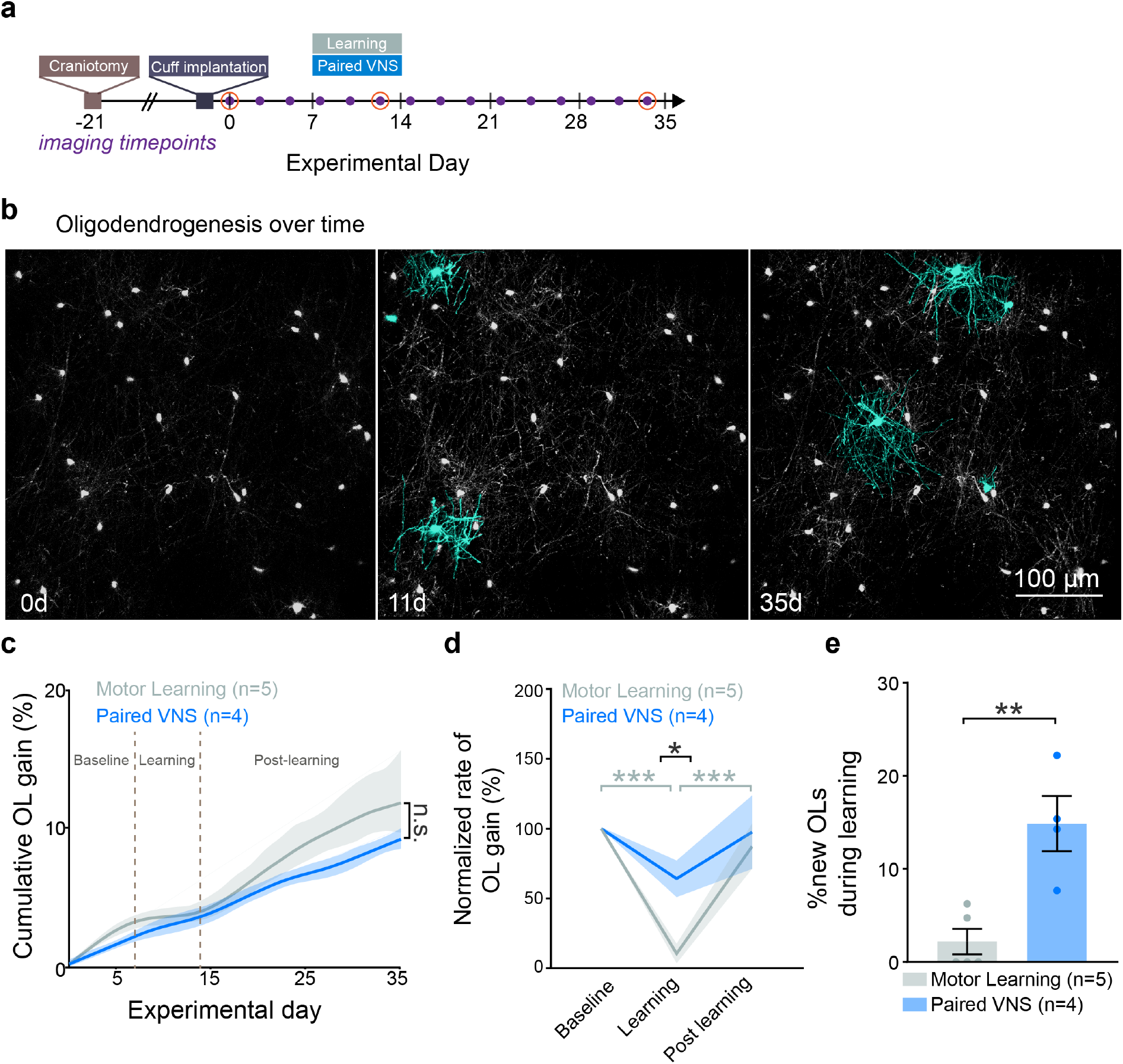
Paired VNS rescues suppression of oligodendrogenesis during motor learning in healthy mice. **a**, Experimental timeline with imaging timepoints. **b**, Representative images of oligodendrogenesis in motor cortex over time with new cells indicated in cyan. **c**, Cumulative oligodendrogenesis (%; relative to baseline) across five weeks is not different between Motor Learning and Paired VNS group (Student’s t-test; p=0.43). **d**, Motor learning modulates oligodendrogenesis rate (p=0.0002), leading to a reduction of oligodendrogenesis rate during learning phase compared to baseline (p=0.0002, Tukey’s HSD). The rate increases in post learning phase (p=0.0009, Tukey’s HSD). VNS rescues this suppression of oligodendrogenesis rate, resulting in a higher oligodendrogenesis rate during learning phase compared to Motor Learning group (p=0.029, Tukey’s HSD). **e**, A larger portion of new cells is generated during learning in Paired VNS group compared to Motor Learning group (Student’s t-test, p=0.0042). Individual dots represent individual mice. Lines and shaded areas in d and e represent the mean ± s.e.m. *p<0.05, **p<0.01, ***p<0.001, ns, not significant; bars and error bars represent the mean ± s.e.m.

**Extended Data Fig. 3.**
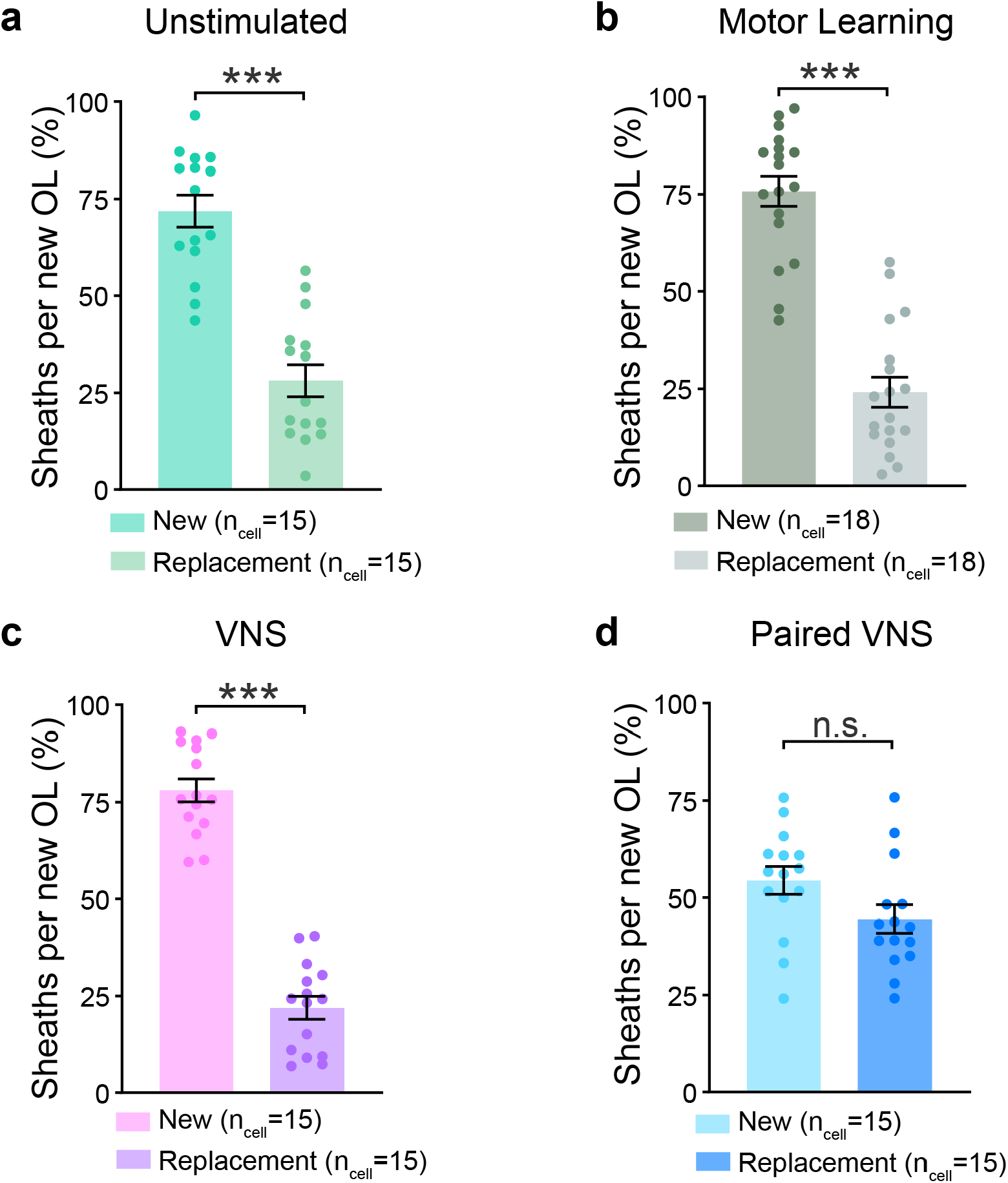
New sheaths from Paired VNS groups exhibit similar chance to myelinate at either unmyelinated or demyelinated area. **a**, New sheaths from Unstimulated mice are more likely to place at previously unmyelinated locations (Paired student’s t-test; p=0.0001). **b**, New sheaths from Motor Learning mice are more likely to place at previously unmyelinated locations (Paired student’s t-test; p<0.0001). **c**, New sheaths from VNS mice are more likely to place at previously unmyelinated locations (Paired student’s t-test; p<0.0001). **d**, New sheaths from Paired VNS mice show equally chance to place either at previously unmyelinated locations or demyelinated locations (Paired student’s t-test; p=0.19).*p<0.05, **p<0.01, ***p<0.001, n.s., not significant; bars and error bars represent the mean ± s.e.m.

**Extended Data Fig. 4.**
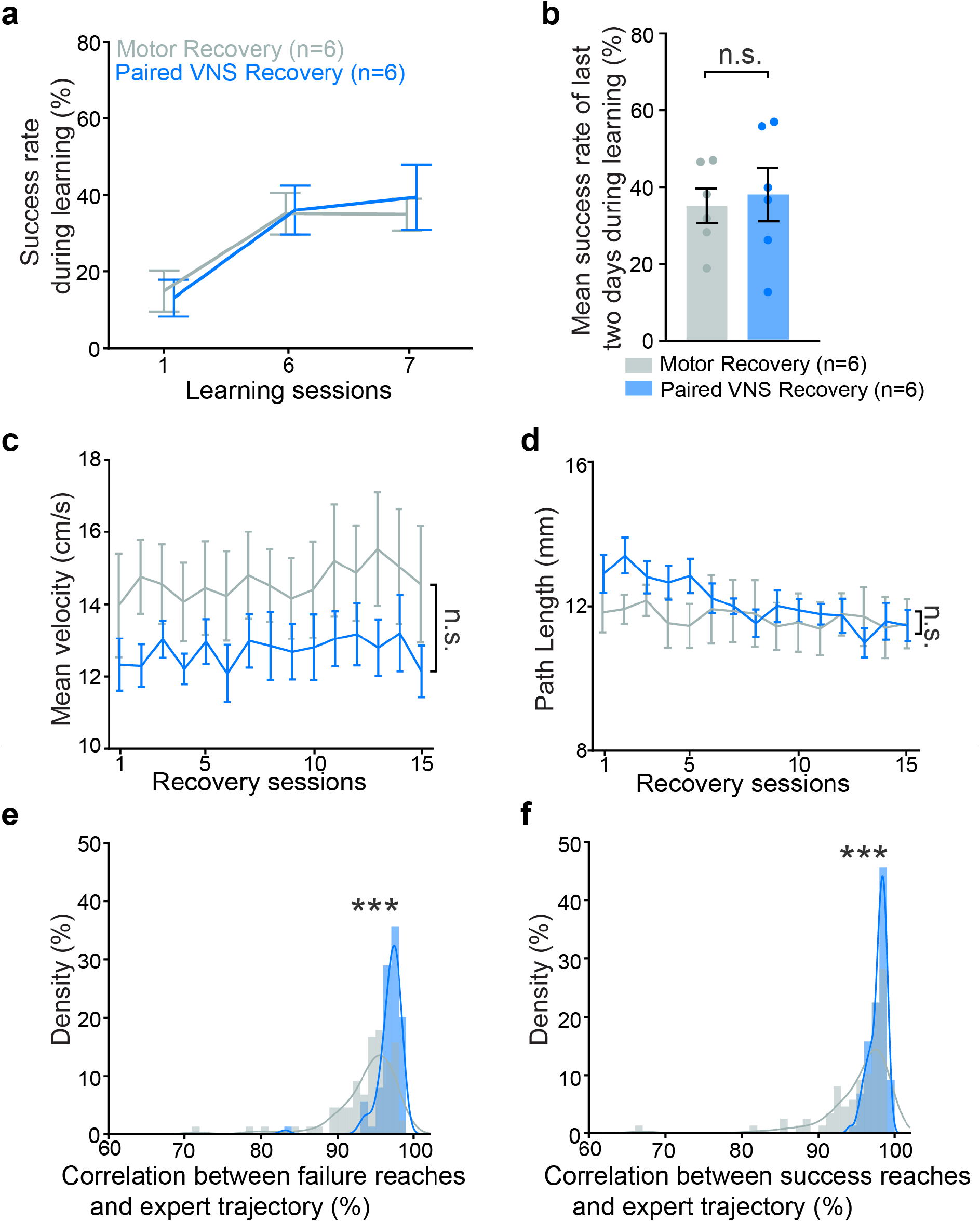
Experimental groups reach a similar level of motor performance prior to demyelination. **a**, Comparison of success rate during learning phase for both groups. Both groups reach similar performance levels before receiving cuprizone diet (REML; p=0.91). **b**, The mean success rate of last two days during learning is similar among both groups (Student’s t-test; t(10)=0.31, p=0.76). **c**, The reach velocity are similar between two groups (REML; F(1)= 1.98, p=0.19). **d**, The length of reach trajectory are similar between two groups (REML; F(1)= 0.35, p=0.57). **e**, The histogram distribution of correlation between failure reaches and expect reaches between both groups is different (Kolmogorov-Smirnov Two-Sample Test; p<0.0001). **f**, The histogram distribution of correlation between success reaches and expect reaches between both groups is different (Kolmogorov-Smirnov Two-Sample Test; p<0.0001).*p<0.05, **p<0.01, ***p<0.001, n.s., not significant; bars and error bars represent the mean ± s.e.m.

