## Supplemental Table for "Paired vagus nerve stimulation drives precise remyelination and motor recovery after myelin loss"

### Data S1 | Statistics results table

*N.B. Statistics results are sorted by figure, and both main figures and supplementary figures are listed in the order they appear in the text.*

#### Table 1.1 | Statistics for Figure 1 and associated extended data (Extended Data Fig. 1)

| FIG. 1 | Measure | Values | N | Statistical test | Significance |
|---|---|---|---|---|---|
| Fig. 1e | Difference in maximum OL loss (%)between two groups (Unstimulated vs. VNS). Bars and error bars represent mean±SEM, individual points represent individual mice | Unstimulated: 77.56±6.13%<br><br>VNS: 64.04±7.86% | Unstimulated: n=5 mice<br><br>VNS: n=5 mice<br><br>Data drawn from three independent batches | Student's t-test<br>t(8)=1.36 | p=0.21 |
| Fig. 1f | Cross-sectional comparison of OL replacement (%) at a 4-week post-cuprizone between Unstimulated and VNS mice. Bars and error bars represent mean±SEM, individual points represent individual mice. | Unstimulated: 55.36±4.32%<br><br>VNS: 71.11±2.79% | Unstimulated: n=5 mice<br><br>VNS: n=5 mice<br><br>Data drawn from three independent batches | Student's t-test:<br><br>t(8)=3.07 | p=0.015<br><br>cohen's d= 2.17 |
| Fig. 1g | Gompertz 3P modelling of cumulative cell gain (%) normalized to maximum cell loss (%) across time in Unstimulated and VNS mice. Shaded area around asymptote represents SEM on curve estimate. | Unstimulated mice:<br>R-square: 0.92<br>Inflection point: 6.14±0.68 days post-cuprizone<br>Growth rate: 0.118±0.015<br>Asymptote: 58.64±2.38%<br><br>VNS mice:<br>R-square: 0.95<br>Inflection point: 11.46±0.64 days post-cuprizone<br>Growth rate: 0.082±0.007<br>Asymptote: 92.27±3.30% | Unstimulated: n=5 mice<br><br>VNS: n=6 mice<br><br>Data drawn from three independent batches | Student's t test comparison curve parameters:<br>Asymptote:<br>t(9)=8.39<br><br>Growth rate:<br>t(9)=2.43<br><br>Inflection point:<br>t(9)=6.00 | Asymptote, p<0.0001<br><br>Growth rate, p=0.038<br><br>Inflection point, p=0.0002 |
| Fig. 1h | Comparison of rate of OL replacement (%) at week1, week2, week3 during post-stimulation phase in cuprizone-treated mice in either the Unstimulated or VNS condition (bars and error bars represent mean±SEM, individual points represent individual mice). | Unstimulated mice:<br>week1: 1.65±0.15%<br>week2: 0.66±0.27%<br>week3: 0.85±0.18%<br><br>VNS mice:<br>week1: 2.89±0.56%<br>week2: 1.81±0.36%<br>week3: 0.93±0.20% | Unstimulated: n=5 mice<br><br>VNS: n=5 mice<br><br>Data drawn from three batch replications | Two-way ANOVA with Bonferroni correction t-test:<br>group*time interaction: F(2,24)=2.01, p=0.16<br><br>time: F(2,24)=10.02, p<0.001<br><br>group: F(1,24)=9.90, p=0.0044 | week1: t(24)=2.74, p=0.034<br><br>week2: t(24)=2.53, p=0.056<br><br>week3: t(24)=0.19, p=0.93 |
| Fig. 1i | Comparison of maximum rate OL replacement calculated between any two imaging time-points during post learning phase between Unstimulated and VNS groups. Boxplots represent Median, min and max, points represent individual mice | Unstimulated: 2.49±0.25%<br><br>VNS: 4.61±1.03% | Unstimulated: n=5 mice<br><br>VNS: n=5 mice<br><br>Data drawn from three separate batches of the same experiment timeline. | Standard deviation is significantly different between two groups, so Welch's test was used.<br><br>F(4,4)=17.63, p=0.017<br><br>t(4.5)=2.00 | p=0.11 |

| Extended Data Fig. 1 | Measure | Values | N | Statistical test | Significance |
|---|---|---|---|---|---|
| Extended Data Fig. 1a | Number of stimuli individual VNS mice received each day across seven days. Bars and error bars represent mean±SEM, individual points represent individual mice. | day1, 16.67±1.31<br>day2, 18.17±1.19<br>day3 20.83±0.87<br>day4, 21.67±0.92<br>day5, 23.5±0.85<br>day6, 25.83±1.14<br>day7, 27.33±1.09 | VNS: n=6 mice<br><br>Data drawn from three independent batches | | |
| Extended Data Fig. 1b | Cumulative OL gain (% relative to baseline) relative to cumulative OL loss (% relative to baseline) for both groups (Unstimulated vs. VNS). | Raw loss (%) and gain (%) at every imaging time-point in every mouse seperated by groups. | Unstimulated: n=5 mice<br><br>VNS: n=6 mice<br><br>Data drawn from three batch replications | Simple linear regression to correlate the amount of gain to the amount of loss for both groups.<br><br>Unstimulated: R-square=0.60, F(1,101)=150.9<br><br>VNS: R-square=0.79, F(1,119)=440.3 | Unstimulated: p<0.0001<br>VNS: p<0.0001 |
| Extended Data Fig. 1c | Comparison of rate of OL replacement (%) at baseline, stimulation phase in cuprizone-treated mice in either the Unstimulated or VNS condition (bars and error bars represent mean±SEM, individual points represent individual mice). | Unstimulated mice:<br>baseline: 2.60±0.39%<br>stimulation: 2.51±0.31%<br><br>VNS mice:<br>baseline: 3.65±1.09%<br>stimulation: 2.71±0.71% | Unstimulated: n=5 mice<br><br>VNS: n=5 mice<br><br>Data drawn from three batch replications | Two-way ANOVA with Bonferroni correction t-test:<br>group*time interaction: F(1,12)=0.37, p=0.55<br><br>time: F(1,12)=0.54, p=0.48<br><br>group: F(1,12)=0.79, p=0.39 | baseline, t(12)=1.06, p=0.62<br><br>stimulation, t(12)=0.20, p>0.99 |
| Extended Data Fig. 1d | Comparison of heart rate change being tested right after cuff implantation surgery between VNS low ΔBR and VNS mice. (bars and error bars represent mean±SEM, individual points represent individual mice ) | VNS low ΔBR: 5.55±1.50%<br><br>VNS: 8.82±2.02% | VNS low ΔBR : n=3 mice<br><br>VNS: n=5 mice<br><br>Data drawn from 3 independent batches | Student's t-test:<br><br>t(6)=1.13 | p=0.303 |
| Extended Data Fig. 1e | Cross-sectional comparison of OL replacement (%) at a 4-week post-cuprizone between Unstimulated, VNS low ΔBR and VNS mice. Bars and error bars represent mean±SEM, individual points represent individual mice. | Unstimulated: 55.36±4.32%<br><br>VNS low ΔBR: 56.65±3.07%<br><br>VNS: 71.11±2.79% | Unstimulated: n=5 mice<br><br>VNS low ΔBR: n=3 mice<br><br>VNS: n=5 mice<br><br>Data drawn from four independent batches | One-way ANOVA:<br>F(2,10)=6.16<br>P=0.018<br><br>Post-hoc analysis using Dunnett with VNS low ΔBR as an control group. | VNS low ΔBR vs. Unstimulated: p=0.96<br><br>VNS ow ΔBR vs. VNS: p=0.046 |

### Data S1 | Statistics results table

*N.B. Statistics results are sorted by figure, and both main figures and supplementary figures are listed in the order they appear in the text.*

#### Table 1.1 | Statistics for Figure 2 and associated extended data (Extended Data Fig. 2)

| Fig. 2 | Measure | Values | N | Statistical test | Significance |
|---|---|---|---|---|---|
| Fig. 2d | Difference in OL loss (%) by 4 weeks post-cuprizone between two groups (Motor Learning vs. Paired VNS). Bars and error bars represent mean±SEM, individual points represent individual mice | Motor Learning: 70.00±5.59%<br><br>Paired VNS: 69.73±7.50% | Motor Learning: n=6 mice<br><br>Paired VNS: n=5 mice<br><br>Data drawn from four separate batches | Student's t-test<br>t(9)=0.03 | p=0.98 |
| Fig. 2e | Cross-sectional comparison of OL replacement (%) at 4-week post-cuprizone between Motor Learning and Paired VNS mice. Bars and error bars represent mean±SEM, individual points represent individual mice. | Motor Learning: 57.06±2.49%<br>Paired VNS: 76.56±6.91% | Motor Learning: n=6 mice<br><br>Paired VNS: n=5 mice<br><br>Data drawn from four separate batches | Student's t-test<br><br>t(9)=2.86 | p=0.019<br>Cohen's d=1.91 |
| Fig.2f | Gompertz 3P modelling of cumulative cell gain (%) normalized to maximum cell loss (%) across time in Motor Learning and Paired VNS mice with a 30 day post-experiment follow-up. Shaded area around asymptote represents SEM on curve estimate. | Motor Learning mice:<br>R-square: 0.95<br>Inflection point: 12.88±0.91<br>Growth rate: 0.073±0.007<br>Asymptote: 77.45±3.75<br><br>Paired VNS mice:<br>R-square: 0.93<br>Inflection point: 14.16±0.62<br>Growth rate: 0.083±0.006<br>Asymptote: 102.17±3.58 | Motor Learning: n=7 mice<br><br>Paired VNS: n=6 mice<br><br>Data drawn from four batch replications | Student's t-test comparison for curve parameters:<br><br>Asymptote: t(11)=4.55<br>Inflection point: t(11)=1.16<br>Growth rate: t(11)=1.03 | Asymptote: p=0.0008<br>Inflection point: p=0.27<br>Growth rate: p=0.33 |
| Fig. 2g | Comparison of rate of OL replacement (%) at week1, week2, week3 during post stimulation phase in cuprizone-treated mice in either the Motor Learning or Paired VNS condition(bars and error bars represent mean±SEM, individual points represent individual mice). | Motor Learning:<br>week1: 2.45±0.36%<br>week2: 1.25±0.27%<br>week3: 1.01±0.14%<br><br>Paired VNS:<br>week1: 3.89±0.41%<br>week2: 2.37±0.38%<br>week3: 1.42±0.08% | Motor Learning: n=6 mice<br><br>Paired VNS: n=5 mice<br><br>Data drawn from four separate batches | Two-way ANOVA with Bonferroni correction t-test:<br>group*time interaction: F(2,27)=1.58, p=0.22<br><br>time: F(2,27)=22.84, p<0.001<br><br>group: F(1,27)=16.80, p<0.001 | week1: t(27)=3.44, p=0.0057<br><br>week2: t(27)=2.68, p=0.037<br><br>week3: t(27)=0.98, p>0.99 |
| Fig 2h | Comparison of maximum rate OL replacement observed between any two imaging time-points during post stimulation phase between Motor Learning and Paired VNS groups. Boxplots represent Median and IQR, values shown as mean±SEM, points represent individual mice | Motor Learning: 3.38±0.42%<br><br>Paired VNS: 7.26±0.53% | Motor Learning: n=6 mice<br><br>Paired VNS: n=5 mice<br><br>Data drawn from four separate batches of the same experiment timeline. | Student's t-test<br><br>t(9)=-5.89 | p=0.0002<br><br>cohen's d= 3.93 |
| Fig 2i | Gompertz 3P modelling of cumulative cell gain (%) normalized to maximum cell loss (%) across time in unstimulated, Motor Learning, VNS and Paired VNS mice. Shaded area around asymptote represents SEM on curve estimate. | Unstimulated mice:<br>R-square: 0.92<br>Inflection point: 6.14±0.68<br>Growth rate: 0.118±0.015<br>Asymptote: 58.64±2.38<br><br>Motor Learning mice:<br>R-square: 0.95<br>Inflection point: 12.88±0.91<br>Growth rate: 0.073±0.007<br>Asymptote: 77.45±3.75<br><br>VNS mice:<br>R-square: 0.95<br>Inflection point: 11.46±0.64<br>Growth rate: 0.082±0.007<br>Asymptote: 92.27±3.30<br><br>Paired VNS mice:<br>R-square: 0.93<br>Inflection point: 14.16±0.62<br>Growth rate: 0.083±0.006<br>Asymptote: 102.17±3.58 | Unstimulated: n=5 mice<br>Motor Learning: n=7 mice<br>VNS: n=6 mice<br>Paired VNS: n=6 mice<br><br>Data drawn from six separate batches of the same experiment timeline. | One-way ANOVA with Tukey's HSD<br><br>asymptote, F(3,20)=27.05, P<0.0001<br><br>Inflection point, F(3,20)=20.99, P<0.0001<br><br>Inflection point, F(3,20)=5.05, P=0.0092 | Asymptote:<br>Unstimulated vs. Motor Learning, p=0.0066<br>unstimulated vs. VNS, p=0.0002<br>unstimulated vs. Paired VNS, p<0.0001<br>Motor Learning vs. VNS, p=0.024<br>Motor Learning vs. Paired VNS, p<0.0001<br>VNS vs. Paired VNS, p=0.22<br><br>Inflection point:<br>Unstimulated vs. Motor Learning, p<0.0001<br>unstimulated vs. VNS, p=0.0004<br>unstimulated vs. Paired VNS, p<0.0001<br>Motor Learning vs. VNS, p=0.57<br>Motor Learning vs. Paired VNS, p=0.50<br>VNS vs. Paired VNS, p=0.070<br><br>Growth rate:<br>Unstimulated vs. Motor Learning, p=0.0066<br>Unstimulated vs. VNS, p=0.045<br>unstimulated vs. Paired VNS, p=0.047<br>Motor Learning vs. VNS, p=0.84<br>Motor Learning vs. Paired VNS, p=0.83<br>VNS vs. Paired VNS, p>0.99 |

| Fig. 2 | Measure | Values | N | Statistical test | Significance |
|---|---|---|---|---|---|
| Fig 2j | Comparison of maximum rate OL replacement observed between any two imaging time-points during post learning phase among four groups. Boxplots represent Median and IQR, values shown as mean±SEM, points represent individual mice | Unstimulated: 2.49±0.25%<br>Motor Learning: 3.38±0.42%<br>VNS: 4.61±1.03%<br>Paired VNS: 7.26±0.53% | Unstimulated: n=5 mice<br>Motor Learning: n=6 mice<br>VNS: n=5 mice<br>Paired VNS: n=5 mice<br><br>Data drawn from six separate batches of the same experiment timeline. | One-way ANOVA with Tukey's HSD<br><br>F(3,17)=11.25, P=0.0003 | Unstimulated vs. Motor Learning, p=0.72<br>unstimulated vs. VNS, p=0.11<br>unstimulated vs. Paired VNS, p=0.0003<br>Motor Learning vs. VNS, p=0.48<br>Motor Learning vs. Paired VNS, p=0.0013<br>VNS vs. Paired VNS, p=0.036 |
| Fig 2k | Cross-sectional comparison of OL replacement (%) at a 4-week post-cuprizone between Unstimulated, Motor Learning, VNS and Paired VNS mice. Bars and error bars represent mean±SEM, individual points represent individual mice. | Unstimulated: 55.36±4.32%<br>Motor Learning: 57.06±2.49%<br>VNS: 71.11±2.79%<br>Paired VNS: 76.56±6.91% | Unstimulated: n=5 mice<br>Motor Learning: n=6 mice<br>VNS: n=5 mice<br>Paired VNS: n=5 mice<br><br>Data drawn from six separate batches of the same experiment timeline. | One-way ANOVA with Tukey's HSD<br><br>F(3,17)=5.76, P=0.0066 | Unstimulated vs. Motor Learning, p=0.99<br>unstimulated vs. VNS, p=0.093<br>unstimulated vs. Paired VNS, p=0.017<br>Motor Learning vs. VNS, p=0.13<br>Motor Learning vs. Paired VNS, p=0.022<br>VNS vs. Paired VNS, p=0.82 |

| Extended Data Fig.2 | Measure | Values | N | Statistical test | Significance |
|---|---|---|---|---|---|
| Extended Data Fig.2c | Comparison of total cumulative OL gain at experimental day 35 between Motor Learning and Paired VNS mice. Solid lines and shaded areas represents mean±SEM. | Motor Learning: 11.60±2.26%<br>Paired VNS: 9.41±0.40% | n=5 Motor Learning mice, with baseline OL counts ranging from 103-160<br><br>n=4 Paired VNS mice with baseline OL counts ranging from 103-147.<br><br>Data drawn from four batches of replicated experiments. | Student's t-test<br>t(7)=0.84 | p=0.43 |
| Extended Data Fig.2d | Normalized rate of OL gain(%; the rate of OL gain is defined as percent increase in population over number of days elapsed; the rate at different phases are normalized to its own rate of OL gain during baseline) baseline (1 week before learning), learning(1 week), and post learning (3 weeks after learning) compared to age-matched controls. Solid lines and shaded areas represents mean±SEM. | Baseline:<br>100% (Motor Learning) and 100% (Paired VNS)<br>Learning:<br>10.67±6.86%(Motor Learning) and 64.17±13.15% (Paired VNS)<br>3w post-learn:<br>87.22±16.28% (Motor Learning) and 97.50±26.30% (Paired VNS) | n=5 Motor Learning mice, with baseline OL counts ranging from 103-160<br><br>n=4 Paired VNS mice with baseline OL counts ranging from 103-147.<br><br>Data drawn from four batches of replicated experiments. | Two-way ANOVA.<br>Group * time interaction:<br>time effects: F(2,21)=13.09, p=0.0002<br>group effects: F(1, 21)=3.81, p=0.065<br><br>Post-hoc analyses using Tukey's HSD<br><br>η2=0.81 | Motor Learning:<br>baseline vs. learning: p=0.0002<br>baseline vs. post learning: p=0.76<br>post-learning vs. learning: p=0.0009<br><br>Paired VNS:<br>baseline vs. learning: p=0.19<br>baseline vs. post learning: p=0.99<br>post-learning vs. learning: p=0.24<br><br>Motor Learning vs. Paired VNS:<br>learning: p=0.029 |
| Extended Data Fig.2e | Percentage of new cells being generated during learning phase over the whole 5 weeks imaging period (baseline + learning + 3w post-learn) relative to age-matched Motor Learning controls. Individual points represent individual mice. Error bars represents mean±SEM. | Motor Learning: 2.20±1.37%<br>Paired VNS: 14.90±2.98% | n=5 Motor Learning mice, with baseline OL counts ranging from 103-160<br><br>n=4 Paired VNS mice with baseline OL counts ranging from 103-147.<br><br>Data drawn from four batches of replicated experiments. | Student's t-test<br>t(7)=4.18 | p=0.0042<br>Cohen's d=3.16 |

### Data S1 | Statistics results table

*N.B. Statistics results are sorted by figure, and both main figures and supplementary figures are listed in the order they appear in the text.*

#### Table 1.3 | Statistics for Figure 3 and associated extended data (Extended Data Fig.3)

| FIG. 3 | Measure | Values | N | Statistical test | Significance |
|---|---|---|---|---|---|
| Fig 3b | Comparison of number of new sheaths from newly generated cells among unstimulated, Motor Learning, VNS and paired VNS groups (bars and error bars represent mean±SEM; individual points represent individual oligodendrocytes) | Unstimulated: 48.60±3.71<br>Motor Learning: 47.43±2.83<br>VNS: 49.80±2.60<br>paired VNS: 52.82±1.79 | Unstimulated: n=10 cells from 5 mice<br>Motor Learning: n=14 cells from 6 mice<br>VNS: n=10 cells from 5 mice<br>Paired VNS: n=11 cells from 5 mice<br>Data drawn from seven independent batches | One-way ANOVA:<br>F(3,41)=0.71 | p=0.55 |
| Fig 3c | Comparison of individual sheath length of new oligodendrocyte from unstimulated, Motor Learning, VNS and Paired VNS mice. Lines represent mean±SEM, points represent individual sheaths | Unstimulate, sheath length: 60.47±1.15 µm<br>Motor Learning, sheath length: 60.72±0.96 µm<br>VNS, sheath length: 58.32±1.18 µm<br>Paired VNS, sheath length: 58.07±1.0.97 µm | Unstimulated: n=447 sheaths<br>Motor Learning: n=655 sheaths<br>VNS: n=446 sheaths<br>Paired VNS: n=585 sheaths<br>Data drawn from seven independent batches | Distribution of sheath lengths violated normality in all four groups: Unstimulated (Shapiro-Wilk, W=0.92, p<0.0001), Motor Learning (Shapiro-Wilk, W=0.94, p<0.0001), VNS (Shapiro-Wilk, W=0.92, p<0.0001) and Paired VNS mice (W=0.94, p<0.0001).<br>Kruskal-Wallis test: statistic is 6.83 | p=0.077 |
| Fig 3d | Comparison of total projected replacement of baseline sheath number by four weeks post-cuprizone among all four groups. Bars and error bars represent mean±SEM, individual points represent individual mice. | Unstimluated mice: -30.46±6.47%<br>Motor Learning mice: -26.72±3.82%<br>VNS mice: -10.08±3.44%<br>Paired VNS mice: -3.16±7.29% | n=5 stimulated mice<br>n=6 Motor Learning mice<br>n=5 VNS mice<br>n=5 paired VNS mice.<br>Data are drawn from seven experimental batches | One-way ANOVA with Tukey's HSD:<br>F(3,17)=5.80, P=0.0064 | Unstimulated vs. Motor Learning, p=0.96<br>unstimulated vs. VNS, p=0.079<br>unstimulated vs. paired VNS, p=0.013<br>Motor Learning vs. VNS, p=0.15<br>Motor Learning vs. paired VNS, p=0.027<br>VNS vs. Paired VNS, p=0.81 |
| Fig 3d | Comparison of total projected replacement of baseline sheath number by four weeks post-cuprizone to baseline. Bars and error bars represent mean±SEM, individual points represent individual mice. | Unstimluated mice: -30.46±6.47%<br>Motor Learning mice: -26.72±3.82%<br>VNS mice: -10.08±3.44%<br>Paired VNS mice: -3.16±7.29% | n=5 stimulated mice<br>n=6 Motor Learning mice<br>n=5 VNS mice<br>n=5 paired VNS mice.<br>Data are drawn from seven experimental batches | One sample t-test with control set as 0 | unstimulated: p=0.0094<br>Motor Learning: p=0.0009<br>VNS: p=0.043<br>Paired VNS: p=0.69 |
| Fig 3f | Comparison of proportion of remyelinating sheaths from newly generated cells post-cuprizone between all four groups (bars and error bars represent mean±SEM; individual points represent individual oligodendrocytes) | Unstimluated mice: 28.16±4.16%<br>Motor Learning mice: 24.19±3.88%<br>VNS mice: 21.97±2.98%<br>Paired VNS mice: 44.58±3.63% | Unstimulated, n=15 cells from 5 mice<br>Motor Learning, n=18 cells from 6 mice<br>VNS, n=15 cells from 5 mice<br>Paired VN, n=15 cells from 5 mice<br>Data are drawn from seven experimental batches | One-way ANOVA with Tukey's HSD:<br>F(3,59)=7.32, P=0.0003 | Unstimulated vs. Motor Learning, p=0.87<br>unstimulated vs. VNS, p=0.66<br>unstimulated vs. paired VNS, p=0.012<br>Motor Learning vs. VNS, p=0.97<br>Motor Learning vs. paired VNS, p=0.0012<br>VNS vs. Paired VNS, p=0.0005 |

| Extended Data Fig.3 | Measure | Values | N | Statistical test | Significance |
|---|---|---|---|---|---|
| Extended Data Fig.3a | Comparison of protpration of newly generated sheaths from newly generated cells that remyelination vs. remodel in Unstimulated mice. Lines represent mean±SEM, points represent individual cells | % sheaths remyelinating: 28.16±4.16<br><br>% sheaths remodeling: 71.84±4.16 | n=15 cells<br><br>Data drawn from three independent batches | Paired student's t-test:<br>t(14)= 5.25 | p=0.0001<br><br>cohen's d=1.98 |
| Extended Data Fig.3b | Comparison of protpration of newly generated sheaths from newly generated cells that remyelination vs. remodel in Motor Learning mice. Lines represent mean±SEM, points represent individual cells | % sheaths remyelinating: 24.19±3.88<br><br>% sheaths remodeling: 75.81±3.88 | n=18 cells<br><br>Data drawn from four independent batches | Paired student's t-test:<br>t(17)= 6.64 | p<0.0001<br><br>cohen's d=2.21 |
| Extended Data Fig.3c | Comparison of protpration of newly generated sheaths from newly generated cells that remyelination vs. remodel in VNS mice. Lines represent mean±SEM, points represent individual cells | % sheaths remyelinating: 21.97±2.98<br><br>% sheaths remodeling: 78.03±2.98 | n=15 cells<br><br>Data drawn from three independent batches | Paired student's t-test:<br>t(14)=9.40 | p<0.0001<br><br>cohen's d=3.55 |
| Extended Data Fig.3d | Comparison of protpration of newly generated sheaths from newly generated cells that remyelination vs. remodel in Paired VNS mice. Lines represent mean±SEM, points represent individual cells | % sheaths remyelinating: 44.58±3.63<br><br>% sheaths remodeling: 54.43±3.63 | n=15 cells<br><br>Data drawn from two independent batches | Paired student's t-test:<br>t(14)=1.38 | p=0.19 |

### Data S1 | Statistics results table

*N.B. Statistics results are sorted by figure, and both main figures and supplementary figures are listed in the order they appear in the text.*

#### Table1.4 | Statistics for Figure 4

| FIG. 4 | Measure | Values | N | Statistical test | Significance |
|---|---|---|---|---|---|
| Fig 4e | Comparison of persistent sheath number by different categories ('Survived' and 'Replaced')between Leanring alone and Paired VNS groups. Bars and error bars represent mean±SEM, individual points represent individual mice. | Survived:<br>Motor Learning: 33.50±5.57<br>Paired VNS: 46.60±11.19<br><br>Replaced:<br>Motor Learning: 22.50±3.10<br>Paired VNS: 40.40±5.54 | Motor Learning: n=6 mice<br>Paired VNS: 5 mice<br>Data drawn from four independent batches | Multiple t test<br>Survived: t(9)=1.11<br>Replaced: t(9)=2.96 | Survived:<br>adjusted p=0.16<br><br>Replaced:<br>adjusted p=0.017<br>cohen's d=1.97 |
| Fig 4f | Comparison of persistent sheath number within selected field of view between Motor Learning and Paired VNS groups. Bars and error bars represent mean±SEM, individual points represent individual mice. | Motor Learning: 56.00±5.61<br><br>Paired VNS: 87.00±12.17 | Motor Learning: n=6 mice<br>Paired VNS: 5 mice<br>Data drawn from four independent batches | Student's t test:<br>t(9)=2.46 | p=0.036<br>cohen's d=1.64 |
| Fig 4g | Comparison of myelin pettern similarity caculated as the persistent sheath number as a ratio of total sheath number (non restored+ persistent+de novo) between Motor Learning and Paired VNS groups. Bars and error bars represent mean±SEM, individual points represent individual mice. | Motor Learning: 39.33±1.56%<br><br>Paired VNS: 48.99±4.01% | Motor Learning: n=6 mice<br>Paired VNS: 5 mice<br>Data drawn from four independent batches | Student's t test:<br>t(9)=2.41 | p=0.039<br>cohen's d=1.61 |

### Data S1 | Statistics results table

*N.B. Statistics results are sorted by figure, and both main figures and supplementary figures are listed in the order they appear in the text.*

#### Table 1.5 | Statistics for Figure 5 and associated extended data (Extended Data Fig.4 )

| FIG 5 | Measure | Values | N | Statistical test | Significance |
|---|---|---|---|---|---|
| Fig. 5b | Comparison of mouse-by-mouse performance in reaching success rate between pre-cuprizone and first three days of post-cuprizone in Motor Recovery and Paired VNS Recovery group. Data shown per individual mouse. | Motor Recovery:<br>Pre cuprizone: 35.14±4.48%<br>Post cuprizone: 24.56±4.03%<br><br>Paired VNS Recovery:<br>Pre cuprizone: 37.75±6.99%<br>Post cuprizone: 44.41±4.49% | Motor Recovery: n=6 mice<br><br>Paired VNS Recovery: n=6 mice<br><br>Data drawn from two batch replications | Paired student's t-test<br><br>Motor Recovery: t(5)= 2.73<br>Paired VNS Recovery: t(5)= 1.02 | Motor Recovery: p= 0.041, cohen's d= 1.01<br>Paired VNS Recovery: p= 0.36 |
| Fig. 5b | Comparison of reaching success rate between Motor Recovery and Paired VNS Recovery group during first three retraining days of post-cuprizone phase . Data shown per individual mouse. | Motor Recovery: 24.56±4.03%<br><br>Paired VNS Recovery: 44.41±4.49% | Motor Recovery: n=6 mice<br><br>Paired VNS Recovery: n=6 mice<br><br>Data drawn from two batch replications | Unpaired Student's t-test<br><br>t(10)= 3.29 | p= 0.0081<br>cohen's d= 2.08 |
| Fig. 5c | Comparison of reaching success rates (% of all attempts that are successful) across all fifteen retraining sessions between Motor Recovery and Paired VNS Recovery groups (points and error bars represent mean±SEM per day per group) | Motor Recovery: 29.17±4.17%<br><br>Paired VNS Recovery: 42.99±3.19% | Motor Recovery: n=6 mice<br><br>Paired VNS Recovery: n=6 mice<br><br>Data drawn from two batch replications | Restricted Maximum Likelihood model (REML) to predict success rate (%) with full factorial model of Day (Retraining day 1-15), Group (Motor Recovery and Paired VNS Recovery) and Day*Group, and with random variable of Mouse nested within Batch.<br><br>Effects of days: F(14)=0.79, p=0.67<br>Effects of groups: F(1)=5.89, p=0.036<br>Interaction Effects: F(14)=1.04, p=0.42 | Benjamini and Hochberg post hoc (false discovery rate(FDR)= 10%)<br>Motor Recovery vs. Paired VNS Recovery:<br>day 2: adjusted p= 0.015<br>day 3: adjusted p= 0.015<br>day 9: adjusted p= 0.066<br>day 10: adjusted p= 0.099<br>day 11: adjusted p= 0.090<br>day 12: adjusted p= 0.090 |
| Fig. 5d | Comparison of minimum reaching success rate between Motor Recovery and Paired VNS Recovery group during the whole retraining sessions . Data shown per individual mouse. | Motor Recovery: 16.87±4.22%<br><br>Paired VNS Recovery: 27.80±2.48% | Motor Recovery: n=6 mice<br><br>Paired VNS Recovery: n=6 mice<br><br>Data drawn from two batch replications | Unpaired Student's t-test<br><br>t(10)= 2.33 | p=0.049<br><br>cohen's d=1.47 |
| Fig. 5f | Comparison of different types of failure reaches between Motor Recovery and Paired VNS Recovery group during the retraining sessions . Data shown per individual mouse. | Motor Recovery:<br>Reach: 44.62±3.68%<br>Grasp: 51.52±3.40%<br>Retrieval: 3.86±0.96%<br><br>Paired VNS Recovery:<br>Reach: 20.31±2.46%<br>Grasp: 74.45±2.62%<br>Retrieval: 5.24±1.30% | Motor Recovery: n=6 mice<br><br>Paired VNS Recovery: n=6 mice<br><br>Data drawn from two batch replications | Multiple student's t test:<br>Reach: t(10)=5.49<br>Grasp: t(10)=5.35<br>Retrieval: t(10)=0.85 | Reach: p=0.0003, cohen's d=3.47<br>Grasp: p=0.0003, cohen's d=3.83<br>Retrieval: p=0.42 |
| Fig. 5h | Comparison of reach consistency within session across all fifteen retraining sessions between Motor Recovery and Paired VNS Recovery groups (points and error bars represent mean±SEM per day per group) | Motor Recovery: 91.74±1.58%<br><br>Paired VNS Recovery: 95.64±0.46% | Motor Recovery: n=6 mice<br><br>Paired VNS Recovery: n=6 mice<br><br>Data drawn from two batch replications | Restricted Maximum Likelihood model (REML) to predict within session reach consistency (%) with full factorial model of Day (Retraining day 1-15), Group (Motor Recovery and Paired VNS Recovery) and Day*Group, and with random variable of Mouse nested within Batch.<br><br>Effects of days: F(14)=1.51, p=0.12<br>Effects of groups: F(1)=5.58, p=0.040<br>Interaction Effects: F(14)=0.78, p=0.69<br><br>R-square = 0.65 | Benjamini and Hochberg post hoc (false discovery rate(FDR)= 10%)<br>Motor Recovery vs. Paired VNS Recovery:<br>day 2: adjusted p= 0.046<br>day 5: adjusted p= 0.046<br>day 7: adjusted p= 0.056 |
| Fig. 5i | Comparison of percentage of expert reaches across all fifteen retraining sessions between Motor Recovery and Paired VNS Recovery groups (points and error bars represent mean±SEM per day per group) | Motor Recovery: 71.59±6.85%<br><br>Paired VNS Recovery: 88.34±2.10% | Motor Recovery: n=6 mice<br><br>Paired VNS Recovery: n=6 mice<br><br>Data drawn from two batch replications | Restricted Maximum Likelihood model (REML) to predict expect reaches (%) with full factorial model of Day (Retraining day 1-15), Group (Motor Recovery and Paired VNS Recovery) and Day*Group, and with random variable of Mouse nested within Batch.<br><br>Effects of days: F(14)=2.91, p=0.0007<br>Effects of groups: F(1)=5.47, p=0.041<br>Interaction Effects: F(14)=0.87, p=0.59<br><br>R-square = 0.77 | Benjamini and Hochberg post hoc (false discovery rate(FDR)= 10%)<br>Motor Recovery vs. Paired VNS Recovery:<br>day 2: adjusted p= 0.039<br>day 3: adjusted p= 0.054<br>day 4: adjusted p= 0.045<br>day 5: adjusted p= 0.094<br>day 6: adjusted p= 0.039<br>day 7: adjusted p= 0.090<br>day 8: adjusted p= 0.090<br>day 12: adjusted p= 0.086<br>day 13: adjusted p= 0.094 |
| Fig. 5j | Comparison of distribution of correlation between failure reaches ad expert trajectory across all fifteen retraining sessions between Motor Recovery and Paired VNS Recovery groups | median value:<br><br>Motor Recovery: 94.83%<br><br>Paired VNS Recovery: 97.84% | Motor Recovery: n=6 mice<br><br>Paired VNS Recovery: n=6 mice<br><br>Data drawn from two batch replications | Kolmogorov-Smirnov two-sample test | p<0.0001 |

| Extended Data Fig.4 | Measure | Values | N | Statistical test | Significance |
|---|---|---|---|---|---|
| Extended Data Fig.4a | Comparison of reaching success rates (% of all attempts that are successful) across during learning (prior to cuprizone treatment) between Motor Recovery and Paired VNS Recovery groups (points and error bars represent mean±SEM per day per group) | Motor Recovery: 28.30±4.19%<br><br>Paired VNS Recovery: 29.53±5.58% | Motor Recovery: n=6 mice<br><br>Paired VNS Recovery: n=6 mice<br><br>Data drawn from two batch replications | Restricted Maximum Likelihood model (REML) to predict success rate (%) with full factorial model of Day (Retraining day 1-15), Group (Motor Recovery and Paired VNS Recovery) and Day*Group, and with random variable of Mouse nested within Batch.<br><br>Effects of days: $F(2)=20.22$, $p<0.0001$<br>Effects of groups: $F(1)=0.031$, $p=0.86$<br>Interaction Effects: $F(14)=0.31$, $p=0.74$<br><br>R-square = 0.79 | |
| Extended Data Fig.4b | Comparison of averaged reaching success rate of last two days during learning phase between Motor Recovery and Paired VNS Recovery group. Data shown per individual mouse. | Motor Recovery: 35.14±4.48%<br><br>Paired VNS Recovery: 37.75±6.99% | Motor Recovery: n=6 mice<br><br>Paired VNS Recovery: n=6 mice<br><br>Data drawn from two batch replications | Unpaired Student's t-test<br><br>$t(10)= 0.31$ | $p=0.76$ |
| Extended Data Fig.4c | Comparison of reach velocity across all fifteen retraining sessions between Motor Recovery and Paired VNS Recovery groups (points and error bars represent mean±SEM per day per group) | Motor Recovery: 14.62±1.20 cm/s<br><br>Paired VNS Recovery: 12.72±0.62 cm/s | Motor Recovery: n=6 mice<br><br>Paired VNS Recovery: n=6 mice<br><br>Data drawn from two batch replications | Restricted Maximum Likelihood model (REML) to predict reach velocity with full factorial model of Day (Retraining day 1-15), Group (Motor Recovery and Paired VNS Recovery) and Day*Group, and with random variable of Mouse nested within Batch.<br><br>Effects of days: $F(14)=1.02$, $p=0.44$<br>Effects of groups: $F(1)=1.98$, $p=0.19$<br>Interaction Effects: $F(14)=0.30$, $p=0.99$<br><br>R-square = 0.82 | |
| Extended Data Fig.4d | Comparison of reach pathlength all fifteen retraining sessions between Motor Recovery and Paired VNS Recovery groups (points and error bars represent mean±SEM per day per group) | Motor Recovery: 11.69±0.62 mm<br><br>Paired VNS Recovery: 12.13±0.24 mm | Motor Recovery: n=6 mice<br><br>Paired VNS Recovery: n=6 mice<br><br>Data drawn from two batch replications | Restricted Maximum Likelihood model (REML) to predict reach length with full factorial model of Day (Retraining day 1-15), Group (Motor Recovery and Paired VNS Recovery) and Day*Group, and with random variable of Mouse nested within Batch.<br><br>Effects of days: $F(14)=2.19$, $p=0.011$<br>Effects of groups: $F(1)=0.42$, $p=0.53$<br>Interaction Effects: $F(14)=1.40$, $p=0.16$<br><br>R-square = 0.68 | |
| Extended Data Fig.4e | Comparison of distribution of correlation between failure reaches ad expert trajectory across all fifteen retraining sessions between Motor Recovery and Paired VNS Recovery groups | median value:<br><br>Motor Recovery: 94.83%<br><br>Paired VNS Recovery: 97.84% | Motor Recovery: n=6 mice<br><br>Paired VNS Recovery: n=6 mice<br><br>Data drawn from two batch replications | Kolmogorov-Smirnov two-sample test | $p<0.0001$ |
| Extended Data Fig.4f | Comparison of distribution of correlation between success reaches ad expert trajectory across all fifteen retraining sessions between Motor Recovery and Paired VNS Recovery groups | median value:<br><br>Motor Recovery: 97.13%<br><br>Paired VNS Recovery: 98.11% | Motor Recovery: n=6 mice<br><br>Paired VNS Recovery: n=6 mice<br><br>Data drawn from two batch replications | Kolmogorov-Smirnov two-sample test | $p<0.0001$ |

### Data S1 | Statistics results table

*N.B. Statistics results are sorted by figure, and both main figures and supplementary figures are listed in the order they appear in the text.*

#### Table 1.6 | Statistics for Figure 6

| FIG 6 | Measure | Values | N | Statistical test | Significance |
|---|---|---|---|---|---|
| Fig. 6b | Comparison of reaching success rates (% of all attempts that are successful) across all seven retraining days in Motor Learning mice (points and error bars represent mean±SEM per day per group) | 35.06±4.89% | n=6 mice<br><br>data drawn from two batches of training | Restricted Maximum Likelihood model (REML) to predict success rate (%) with full factorial model of Day (learning days 1-7) and with random variable of Mouse nested within Batch.<br><br>Effects of days:F(6)=1.98, p=0.10<br><br>R-square = 0.64 | Post-hoc tests of interest:<br><br>day 1 vs. day 7: p=0.89 |
| Fig. 6c | Comparison of mouse-by-mouse improvement in reaching success rate between days 1 and 7 of retraining phase in Motor Learning mice. Data shown per individual mouse. | day 1: 32.82±5.30%<br><br>day7: 37.97±3.60% | n=6 mice<br><br>data drawn from two batches of training | Paired student's t-test between day 1 and day 7<br><br>t(5)= 0.87 | p=0.42 |
| Fig. 6d | Comparison of reaching success rates (% of all attempts that are successful) across all seven retraining days in Paired VNS mice (points and error bars represent mean±SEM per day per group) | 39.64±4.23% | n=8 mice<br><br>data drawn from two batches of training | Restricted Maximum Likelihood model (REML) to predict success rate (%) with full factorial model of Day (learning days 1-7) and with random variable of Mouse nested within Batch.<br><br>Effects of days: F(6)=3.35, p=0.0087<br><br>R-square = 0.82 | Post-hoc tests of interest:<br><br>day 1 vs. day 7: p=0.006 |
| Fig. 6e | Comparison of mouse-by-mouse improvement in reaching success rate between days 1 and 7 of retraining phase in Paired VNS mice. Data shown per individual mouse. | day 1: 34.94±5.19%<br><br>day7: 49.20±5.25% | n=8 mice<br><br>data drawn from two batches of training | Paired student's t-test between day 1 and day 7<br><br>t(7)= 3.61 | p=0.0086<br><br>cohen's d=0.97 |
| Fig. 6f | Correlation between Mean success% during retraining and OL number restoration by 4 weeks post cuprizone | | n=8 mice, with 4 mice from Motor Learning group and 4 mice from Paired VNS group. | Standard least squares regression<br>$R^2$=0.001 | p=0.95 |
| Fig. 6g | Correlation between Mean success% during retraining and sheath number restoration by 4 weeks post cuprizone | | n=8 mice, with 4 mice from Motor Learning group and 4 mice from Paired VNS group. | Standard least squares regression<br>$R^2$=0.14 | p=0.36 |
| Fig. 6h | Correlation between Mean success% during retraining and myelin pattern restoration by 4 weeks post cuprizone | | n=8 mice, with 4 mice from Motor Learning group and 4 mice from Paired VNS group. | Standard least squares regression<br>$R^2$=0.65 | p=0.016 |
